# Using deep learning predictions reveals a large number of register errors in PDB deposits

**DOI:** 10.1101/2024.07.19.604304

**Authors:** Filomeno Sánchez Rodríguez, Adam J. Simpkin, Grzegorz Chojnowski, Ronan M. Keegan, Daniel J. Rigden

**Affiliations:** Institute of Systems, Molecular and Integrative Biology, University of Liverpool, Liverpool L69 7ZB, England; Life Science, Diamond Light Source, Harwell Science and Innovation Campus, Didcot, Oxfordshire OX11 0DE, England; Department of Chemistry, York Structural Biology Laboratory, University of York, York, UK; European Molecular Biology Laboratory, Hamburg Unit, Notkestrasse 85, 22607 Hamburg, Germany; UKRI-STFC, Rutherford Appleton Laboratory, Research Complex at Harwell, Didcot OX11 0FA, England

## Abstract

The accuracy of the information in the Protein Data Bank (PDB) is of great importance for the myriad downstream applications that make use of protein structural information. Despite best efforts, the occasional introduction of errors is inevitable, especially where the experimental data are of limited resolution. We have previously established a novel protein structure validation approach based on spotting inconsistencies between the residue contacts and distances observed in a structural model and those computationally predicted by methods such as AlphaFold 2. It is particularly well-suited to the detection of register errors. Importantly, the new approach is orthogonal to traditional methods based on stereochemistry or map-model agreement, and is resolution-independent. Here we identify thousands of likely register errors by scanning 3-5Å resolution structures in the PDB. Unlike most methods, application of our approach yields suggested corrections to the register of affected regions which we show, even by limited implementation, lead to improved refinement statistics in the vast majority of cases. A few limitations and confounding factors such as fold-switching proteins are characterised, but we expect our approach to have broad application in spotting potential issues in current accessions and, through its implementation and distribution in CCP4, helping ensure the accuracy of future deposits.

## 1. Introduction

For more than five decades the Protein Data Bank (PDB; (wwPDB consortium, 2019)) has been collecting experimentally determined macromolecular structures. Currently with over 200,000 entries, its structures come from macromolecular crystallography (MX), nuclear magnetic resonance (NMR) and, more recently, cryogenic electron microscopy (cryo-EM). The bulk of deposits derive from MX, but cryo-EM is on an accelerating trend (Callaway, 2020). Whatever the method, the experiment typically results in the determination of a model that is considered by the authors to (best) satisfy the observations. However, as with all scientific endeavour, experimental limitations can lead to unavoidable uncertainties and hence, despite the experimentalists’ best efforts, the introduction of errors into the final model.

The 1990s saw a recognition of the need for error detection or structure validation tools and a first generation of methods emerged. These were variously based on geometric and stereochemical properties (e.g. PROCHECK (Laskowski *et al*., 1993) and WHATIF (Vriend, 1990)); consideration of statistics of favoured amino-acid environments (e.g. VERIFY_3D (Lüthy *et al*., 1992)) statistics of interatomic contacts (e.g. ERRAT (Colovos & Yeates, 1993) and DACA (Vriend & Sander, 1993), or a combination of a Cβ–Cβ (or Cα–Cα) potential and solvent-exposure statistics (ProSA (Sippl, 1993)). More recent methods have furthered refined these concepts e.g. MolProbity (Davis *et al*., 2007) and have introduced measures of map-model scoring in programs such as EMRinger (Barad *et al*., 2015), Q-score (Pintilie *et al*., 2020) and SMOC (Joseph *et al*., 2016). The very recent method MEDIC marries the geometric and the map-fitting approaches with special application to cryo-EM structures (Reggiano *et al*., 2023). Of particular note, COOT (Casañal *et al*., 2020) and ISOLDE (Croll, 2018) each now provide sophisticated visualisation of diverse structure validation results at the interactive model building stage of structure determination.

A key principle of structure validation is that the property used for validation should not be included in the target function during structure determination. Basic chemical features such as bond distances and angles therefore have limited validation utility since they are typically restrained to ideal values during structure determination, especially in recognition of the fact that many X-ray crystal structures are under-determined, with fewer observations than model parameters. The danger of failing to maintain a distinction between refined parameters and those used for validation is illustrated by the introduction of Ramachandran plot restraints which have led to the emergence of structures that pass Ramachandran tests for outliers but which more sophisticated statistical analysis shows exhibit highly skewed and implausible Ramachandran plot distributions (Afonine *et al*., 2023). The ingenuity of researchers in finding novel, independent metrics such as CABLAM (Prisant *et al*., 2020) has proved important, but new validation metrics are still very valuable.

One particularly difficult class of error to detect is the sequence register error in which the main chain may be broadly correct, but residues are systematically assigned the identity of a residue a number of amino acids up or down in the sequence. Such errors are particularly easily introduced where the local resolution of the map is lower, and especially where more easily identifiable marker residues such as aromatic amino-acids are absent. The fact that the backbone is often essentially correct in the region of the register error means that some conventional validation scores will struggle to detect a problem, although non-rotameric side chains and steric clashes may sometimes be helpful (Chojnowski, 2022, 2023). By representing residue probabilities at each Cα position in an input using a neural network classifier, the program checkMySequence has contributed significantly to the detection of register errors specifically, in both MX and cryo-EM structures (Chojnowski, 2022, 2023), but its dependence on map quality means that its sensitivity naturally declines at poorer resolution. The development of the DAQ score (Terashi *et al*., 2022) and an associated database (Nakamura *et al*., 2023) are further major contributions to structure validation but that analysis too is dependent on map quality and resolution.

We have recently introduced new methods for protein structure validation (Sánchez Rodríguez *et al*., 2022) based on the compatibility of a structure with the inter-residue distances and contacts predicted by methods such as AlphaFold 2 (Jumper *et al*., 2021). These were first a Support Vector Machine (SVM) classifier which predicts whether a given residue is part of an error of any kind; and secondly a contact map alignment procedure which detects whether the mapping between the observed residue contacts in a structure and the predicted contacts is optimal or whether an alternative alignment would score better, a sign of a possible register error. After specific steps to filter out potential false positives based on either a relative paucity of contacts for the potential error region or excessive structural divergence between the structure and the AF2 model, we found a combination of the SVM and contact map alignment to be very effective in detection of sequence register errors. Notable features of our method are its map-independence, making it insensitive to the resolution of the structure analysed, and its ability to use the contact map alignment step to suggest the correction required to arrive at the right sequence register.

Here we present the results of validating all 3-5Å resolution structures in the PDB, both cryo-EM and MX, with our new method. Even applying stringent criteria, we identify thousands of putative register errors which we further validate by comparison with high-resolution crystal structures and with the orthogonal map-based validation tool checkMySequence (Chojnowski, 2022, 2023). Finally, automated correction of the likely errors results, in most cases, in improved local real-space correlation coefficients, adding further confidence that an error has been identified. While our method remains limited by factors that affect prediction of residue distances and contacts by AF2 such as the depth of the MSA that can be constructed of the target sequence and homologues, and can be misled by rare fold-switching proteins, it is powerful and conceptually independent of structure validation methods hitherto applied to PDB entries (Sali *et al*., 2015; Kleywegt *et al*., 2024). These results should therefore help avoid much misguided research effort based on locally incorrect PDB structures (Gao *et al*., 2023).

## 2. Materials and methods

### 2.1 Selection of protein models deposited in the PDB

The dataset of protein structures used in this study was selected by first retrieving a list of all structures determined using cryo-EM or X-ray crystallography which were deposited up to 5th of April 2022 in the PDB at resolutions between 3.0 and 5.0Å. The resulting 19,310 PDB entries were then split into their constituent chains, leading to the creation of a dataset of 203,533 protein chains. Chains solely formed by nucleotides or ligands were discarded. Additionally, chains with more than 1,000 residues were also discarded as obtaining AlphaFold 2 predictions would be intractable due to hardware limitations. The remaining set of 148,785 protein chains were then clustered according to their sequence identity: protein chains sharing 100% sequence identity were grouped together, resulting in the creation of a final dataset of 30,731 clusters, each having an approximate average of 5 members. To ensure a match between the residue numbering observed in the deposited models and the numbering in the reference sequence deposited in the PDB used to obtain AlphaFold 2 predictions, a pairwise sequence alignment was performed using the BLOSUM62 substitution matrix and the chains in these clusters were renumbered accordingly.

### 2.2 Prediction of contact maps derived from of inter-residue distances obtained using AlphaFold 2

Predictions of inter-residue distances were obtained using AlphaFold 2 for each of the clusters in the dataset created as described in section 2.1. Since all the structures in each cluster have 100% sequence similarity, only one prediction was carried out for each individual cluster. For efficiency, the database search required as part of the AlphaFold 2 run was carried out using MMseqs2 API (Mirdita *et al*., 2019) instead of the original jackhmmer search (Eddy, 2011). The original CASP14 model preset was then used and all parameters were left to their defaults. Using this setup, one predicted model was produced with AlphaFold 2 for each of the clusters in the dataset. Then, the inter-residues distances for this predicted model were taken. These predictions consist of the predicted probabilities that each residue pair in the structure is within a given distance bin. Contact predictions were derived from these inter-residue distance predictions by adding together all probabilities observed for the distance bins up to 8Å for each residue pair. Finally, the top L/2 residue contacts scoring the highest probability values were then taken to form the final predicted contact maps, where L denotes the sequence length of the protein chain.

All of these predictions were carried out on a computing grid where each node was equipped with a twin 16-core Intel Xeon Gold 5218 running at 2.3 GHz, 160 GB of memory and four NVIDIA Tesla V100 chips with 16 GB of video memory each.

### 2.3 Contact map alignment-based model validation

Model validation was carried out for each protein model present in the dataset described in section 2.1 using the contact map alignment step of the conkit-validate pipeline (Sánchez Rodríguez *et al*., 2022). In this pipeline, the contact map predicted by AlphaFold 2 was aligned with the contact map derived from the residue distances observed in the deposited model using map_align (Ovchinnikov *et al*., 2017). This tool introduces and extends gaps as required in order to obtain the optimal alignment between the two input contact maps, which is defined as the alignment where the maximum number of contacts match each other in both maps (contact maximum overlap, CMO). In cases where the maximum overlap between contacts was achieved using a sequence register different to the one observed in the deposited model in at least five consecutive residues, the mismatch was flagged by conkit-validate as a possible sequence register error and the sequence register that achieved the CMO was proposed as a possible fix for the predicted error. Additionally, this pipeline merged consecutive regions flagged as register errors if they were separated by three residues or fewer: it seemed unlikely that nearby predicted errors had independent causes and so counting them once seemed fairer. As a final step, errors predicted with this pipeline were filtered using three criteria established in a previous study (Sánchez Rodríguez *et al*., 2022). These criteria were designed in order to discard predicted errors detected at parts of the model where the contact map misalignment could be caused by reasons other than a register error. These three criteria discarded errors where either the contact maps did not contain sufficient information to produce a reliable alignment, the AlphaFold 2 predictions did not have sufficient confidence, or the predicted model could have been modelled in an alternative conformation different to the one of the deposited model.

### 2.4 Selection of high resolution crystal structure for cross-validation

In order to obtain a set of high-resolution structures that could be used to cross-validate the predicted register errors detected in this study, an advanced search was carried out in the PDB to find entries sharing a 100% sequence identity with each of the clusters in the dataset described in section 2.1. To restrict analysis to higher-resolution, generally more confident structures, the results of this search were then filtered so that only structures determined at least at 2.5Å and using X-ray crystallography were left. In cases where multiple entries meeting these criteria were found, the structure determined at the highest resolution was taken.

### 2.5 Modification of sequence register in models with predicted errors

For the deposited models where a possible sequence register error was detected, a model with the alternative sequence register predicted to be correct was created as follows.

First, for each error detected in the structure, a buffer was added at each side of the predicted error to account for cases where the register error was preceded or followed by a modelling error. The length of this buffer was proportional to the predicted residue shift error, and was calculated as follows:

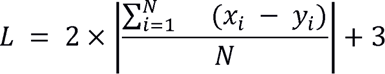

Where L is the length of the buffer, N is the number of residues affected by the predicted error, x is the residue number observed in the deposited structure and y is the number of a corresponding residue predicted to be correct.

In those instances where the buffer regions of two contiguous errors shared any number of residues, the errors were joined and considered as a single error for the purpose of creating a model with the alternative sequence register, but they stayed as separate errors for the rest of the analysis.

Next, a Cα-based sequence-independent structural alignment between the deposited model and the model predicted by AlphaFold 2 was carried out using GESAMT (Krissinel, 2012). Those residues predicted to be part of a sequence register error were then removed from the deposited model, together with their buffers, and replaced with the equivalent set of residues present at the superposed predicted model, which were modelled with the register predicted to be correct. COOT (Emsley & Cowtan, 2004) was then used to perform real-space refinement on each pair of residues where the original deposited model was adjacent to the fragment originating from the predicted model. Further refinement and calculation of quality metrics was then performed on both the deposited model and the newly created model with the alternative sequence register. This was carried out in a different manner depending on the method used to determine the deposited model.

In cases where the structure was originally determined using cryo-EM, REFMAC5 (Murshudov *et al*., 2011) was used to perform 20 cycles of jelly-body refinement both on the original model and the alternative model. Then, the local correlation coefficient between the map and the modified set of residues in both the original and the alternative model was calculated using PHENIX.MAP_MODEL_CC from the PHENIX suite (Liebschner *et al*., 2019).

For MX cases, after correcting the predicted sequence register errors, all the chains present in the asymmetric unit - regardless of whether an error was detected in them or not - were used as input to perform 20 cycles of restrained refinement using REFMAC5, to help fit the modified region to the density. To allow like-for-like comparisons, 20 cycles of restrained refinement using default restraints and automatic weights were also run on the deposited structure. Density-fitness (GitHub - PDB-REDO/density-fitness: Application to calculate the density statistics (RSR, SRSR, RSCCS, EDIAm and OPIA) for x-ray structures), of the PDB-redo suite of programs (Joosten *et al*., 2012), was used to calculate a per-residue Real-Space Correlation Coefficient (RSCC). By iterating through the residues of interest, a mean local RSCC was calculated for the modified and unmodified structures in the region of the predicted register error. For visualising the output MTZ files in Chimera, the MTZ files were converted to MRC files using COOT.

### 2.6 Alternative validation methods

#### 2.6.1 checkMySequence

The map-model compatibility tool checkMySequence (cMS) was applied to all cryo-EM or crystal structures in the set using the procedures as described previously (Chojnowski, 2022, 2023).

#### 2.6.2 Geometry

Unusual geometric features - Ramachandran outliers, side chain rotamer outliers and Cβ distortions were detected with Molprobity (Prisant *et al*., 2020; Davis *et al*., 2007).

#### 2.6.3 DAQ scoring

To further assess the register errors in the cryo-EM structures, the average DAQ score across the residues corresponding to each predicted error was mined from the DAQ score database (Terashi *et al*., 2022) These were compared to the delta PHENIX.MAP_MODEL_CC score between the original and alternative models.

## 3. Results and Discussion

### 3.1 One in six models determined at 3-5Å deposited in the PDB contain a putative register error

Processing of PDB entries at between 3-5Å as outlined in Methods produced a set of 16,662 structures. These were processed using the conkit-validate pipeline, with the purpose of detecting possible register errors. During this analysis, a total of 12,674 possible register errors were found distributed among 2,954 entries (17%). When taking in consideration the fact that some sequences are represented several times in the dataset in the form of homomeric structures or the same protein in different PDB entries, the total number of unique predicted register errors was 4,606. Analysis of the distribution of PDB depositions with at least one predicted error across different resolution bins and deposition years was carried out (Figure 1). Unsurprisingly, predicted errors are more common in lower resolution entries: around 10% of entries at 3.0Å contain a predicted error, rising to around 20% between 3.7Å and 5.0Å. Perhaps less expected is the historical trend towards a larger number of predicted errors recently - only 8% of entries deposited between 1995-2000 contain a predicted error, rising to almost 20% in recent years. However, this presumably reflects the increasing presence of cryo-EM in structure determination more recently and the encouragement its experimentally measured maps offer to lower resolution structural analysis (see below).

**Figure 1.**
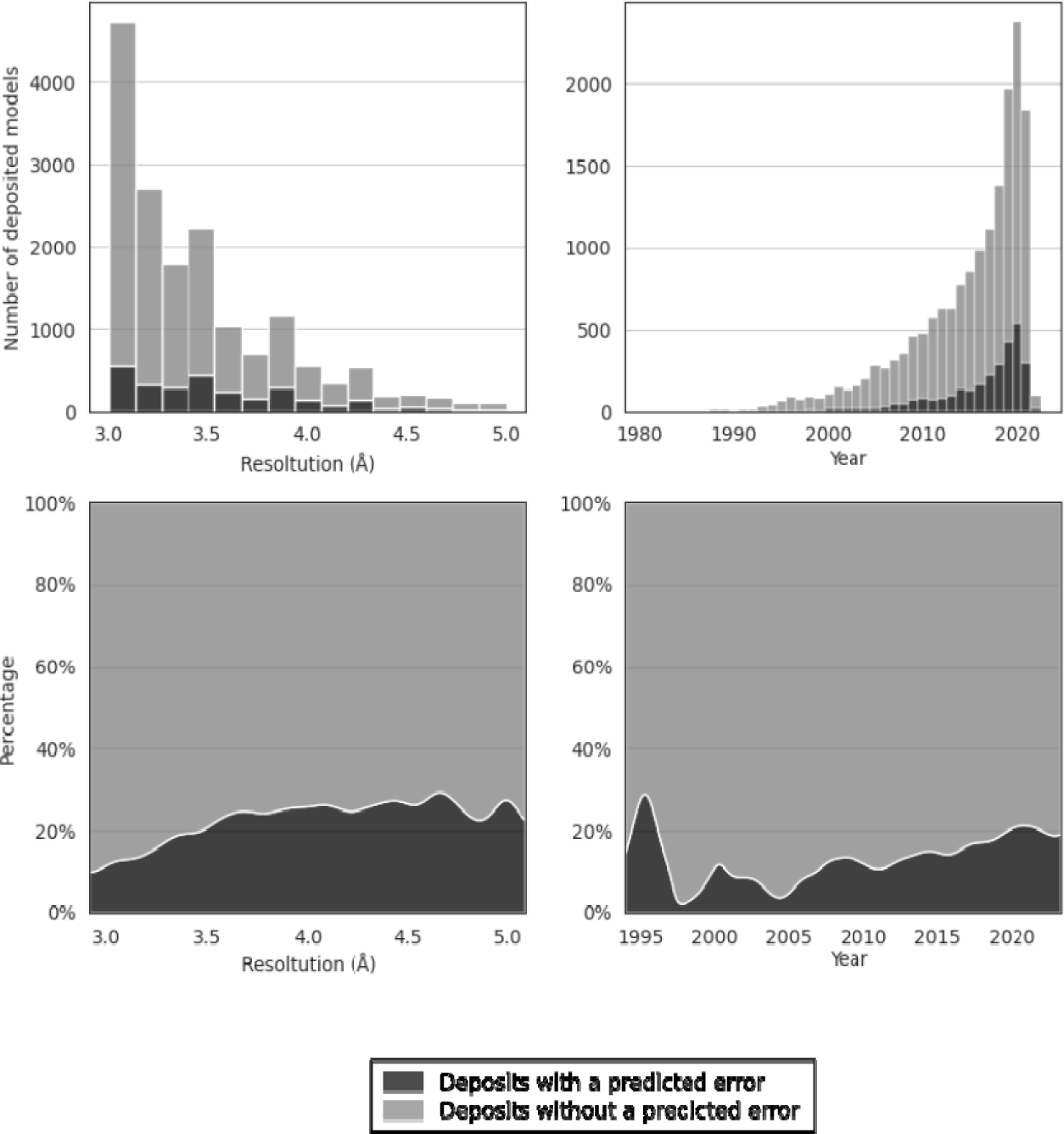
Distribution of models deposited in the PDB where a sequence register error was predicted across different resolution bins (left column) or year of deposition (right column). Plots in the top row depict the stacked count of deposited models without a predicted error (light grey) and the models where an error was predicted (dark grey). Plots in the bottom row show the percentage of the deposited structures where an error was predicted. Depositions before 1995 have been omitted for clarity because the number of entries is small (only 201 in total from 1981-1995). Note that sequence redundancy removal between different PDB entries has not been applied here: some entries represented here will contain the same error.

Further characterisation of the predicted errors was carried out by studying the sequence shift predicted to be required to correct the error, the number of residues affected by the possible error and the fraction of residues present in the overall structure that were found to be part of the error (Figure 2). Most of the predicted errors are small with two thirds affecting 15 residues or fewer. The shifts required to correct predicted errors are also small: two thirds involve a single residue.

**Figure 2.**
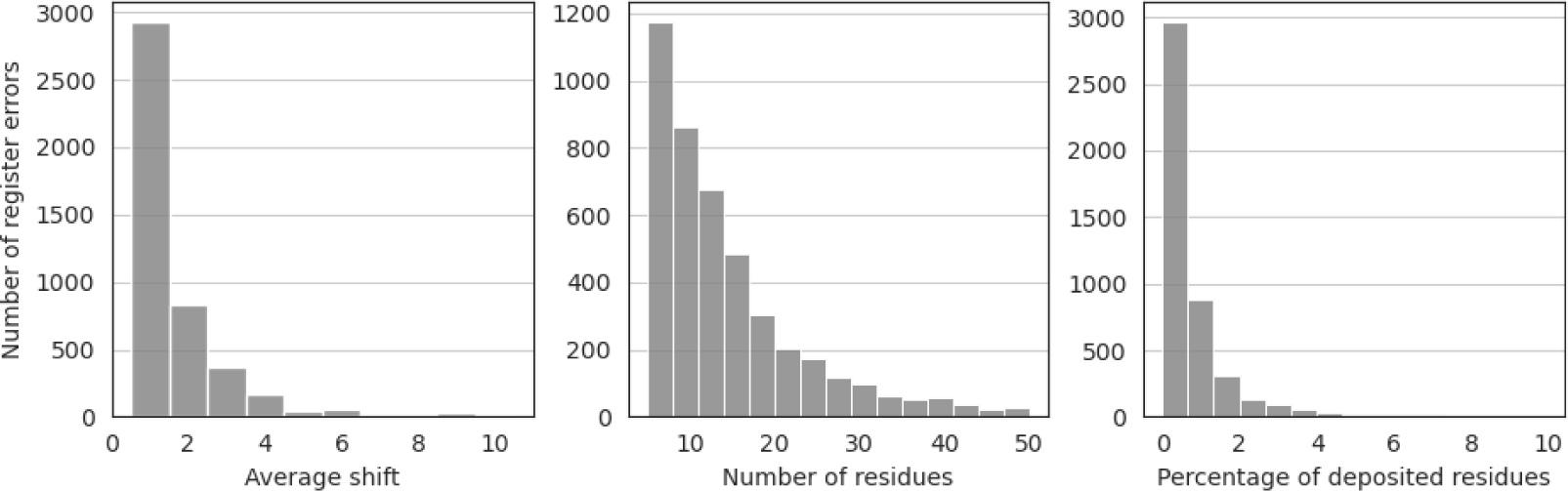
Number of predicted sequence register errors plotted by the predicted residue shift (left), by the number of affected residues (middle) and by the fraction that the affected residues represent compared with the overall structure (right). Errors consisting of a shift of more than 10 residues, affecting more than 50 residues or more than 10% of the residues in the structure have been omitted for clarity (127, 248 and 27 errors respectively). Note that where two errors were separated by three residues or fewer they were combined (see Materials and Methods), meaning where the two errors involved shifts of different numbers of residues, that the average shift for a predicted register error may be non-integral. Note that numbers here are after sequence redundancy removal and represent unique errors within and between PDB entries.

### 3.2 Errors are more common among PDB deposits determined by cryo-EM than by MX

A comparison of the distribution of predicted errors found in structures solved with cryo-EM or MX was carried out. Interestingly, a lower ratio of structures containing errors was observed for MX, regardless of the year of submission or the resolution bin. In contrast with MX error incidence, which was broadly stable over time, the proportion of cryo-EM structures which contain a predicted error was observed to decrease over the recent years, with values approaching those observed among MX structures.

**FIgure 3.**
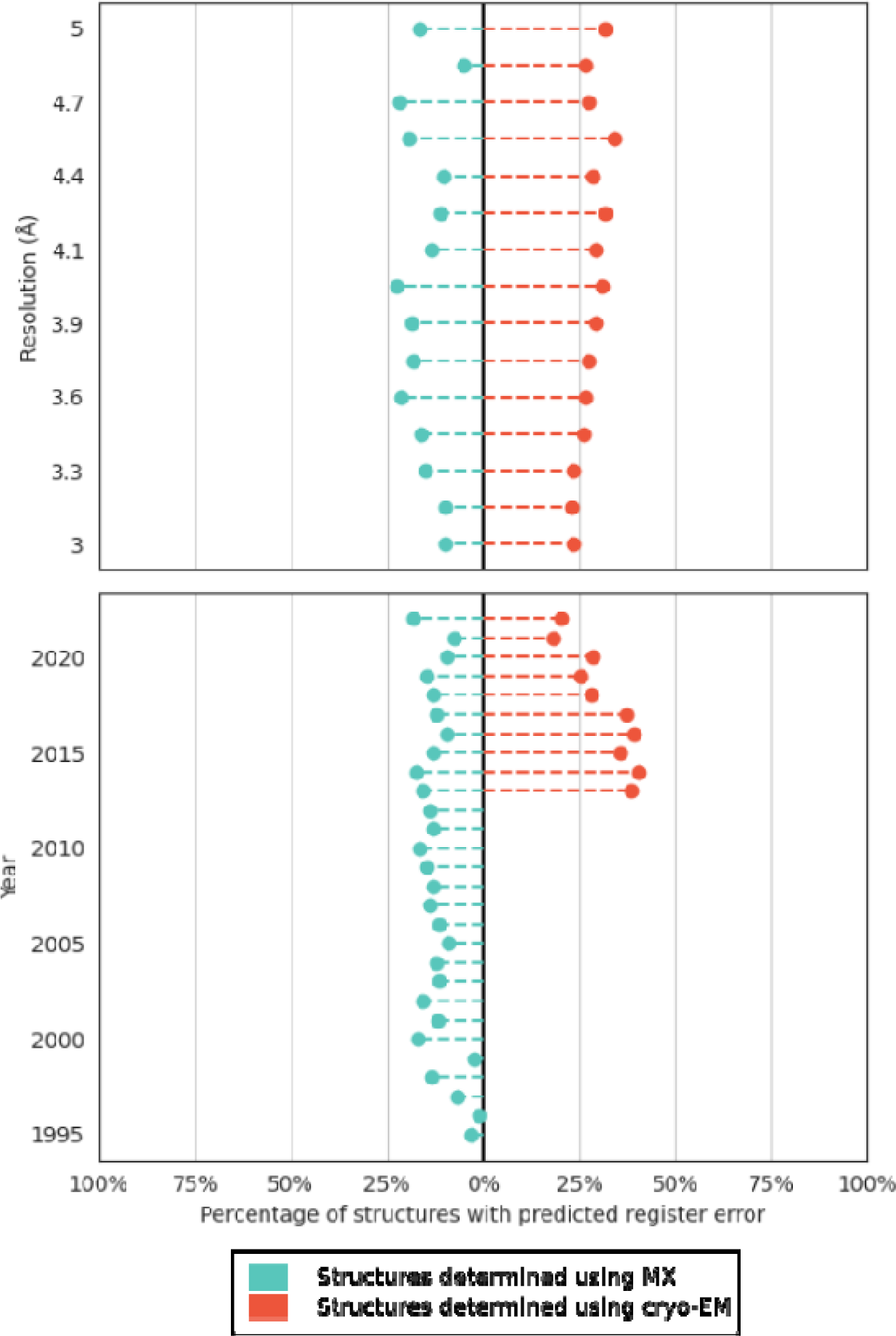
Proportion of structures containing a register error across different resolutions and years. Ratios for structures determined using MX have been coloured in blue while those determined with cryo-EM in red. Data for MX structures deposited before 1995 and cryo-EM structures deposited before 2013 are small in number and have been omitted for clarity (136 and 28 entries respectively).

Breaking down the incidence of cryo-EM and MX predicted errors into resolution bins (Supp Fig 1) suggests that the recent reduction in incidence of predicted registered errors for cryo-EM structures seems to be centred principally on the 3.5A-4.5Å resolution range.

### 3.3 High-resolution crystal structures validate many predicted errors

In order to cross-validate the errors that we found in deposited models, we used the same model validation method to analyse high resolution counterparts of crystal structures from our low-resolution set. We searched for structures with 100% sequence identity, solved using MX at a resolution of at least 2.5Å. In total we found 147 structures that met these criteria and could be used to cross-validate our results: our proposal is that where the later (in 79% of cases), higher-resolution structure does not contain the predicted error found in its lower-resolution predecessor, then the correction of an error with the benefit of better data is the likely explanation. These 147 structures covered 403 unique errors representing, since there were 4606 unique errors in total, approximately 10% of the unique errors (i.e. after sequence redundancy removal) that we found in the original set of depositions. This small set of errors is representative of the full set of errors found in this study in terms of distribution of register error size and shift (Supp Fig 2).

After repeating the validation exercise on the set of 147 structures, no register error was predicted using our approach for 115 of these high-resolution depositions, directly validating (since these structures encompassed 350 predicted errors) 87% of these unique predicted errors (Figure 4).

**Figure 4.**
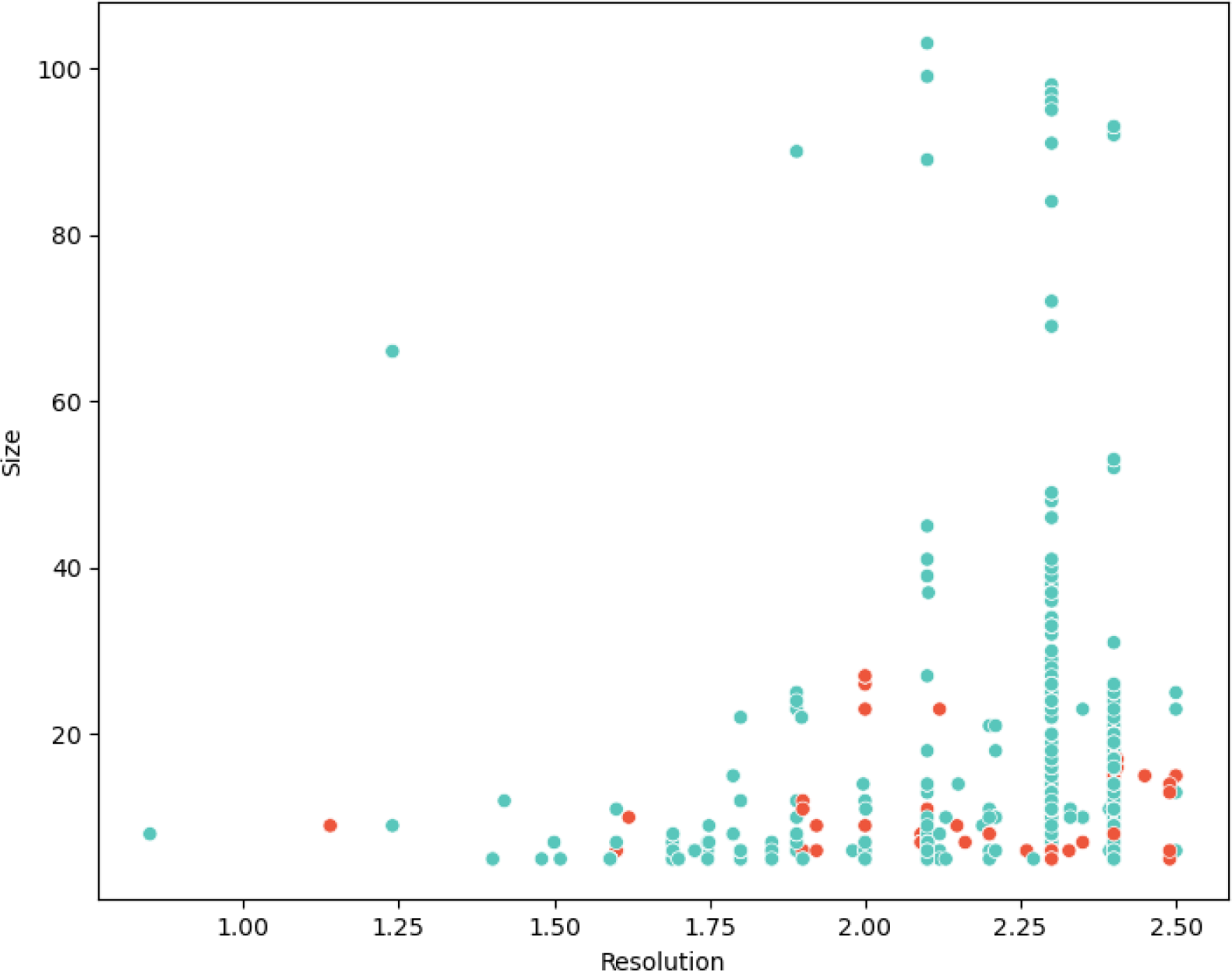
Validation of 403 predicted unique errors by comparison with 147 high-resolution MX structures. Each predicted unique error is plotted according to the resolution of the structure in question and the number of residues affected by the predicted error. Blue points indicate cases where no error is detected in the high resolution crystal structure, red where that is not the case.

Supp Fig 3 shows a typical case where the electron density unambiguously supports the sequence register in the high resolution 1.42 Å structure (of Ragulator complex protein LAMTOR4; PDB code 6B9X, chain D, residues 24-36), thereby validating the prediction of a register error in the lower resolution 3.01 Å structure (PDB code 5YK3, chain I).

Cases where a predicted error is called in the lower resolution structure but also in the higher resolution structure are more surprising where the quoted resolution would generally be considered sufficient to allow unambiguous sequence tracing. The three highest resolution examples were examined in more detail.

The first was a predicted one-residue register shift error spanning eight residues from positions 63-70 in the 3Å MX structure of an *Escherichia coli* FimH complex (PDB 4xob, chain C). The 1.14Å MX structure of the same protein (PDB 4xo9, chain A) was also predicted to have a register error in the same region and had a similar local structure. The AF2 model had a different register in the region - the source of the predicted error in both MX structures - but, unexpectedly, a further high-resolution 1.0 Å structure of (one domain of) FimH (PDB 4x5p, chain A) shared the AF2 register. The electron density for each high-resolution structure unambiguously supports the atomic interpretation (Supp Fig 4a) suggesting that the region is genuinely structurally ambivalent. A possible explanation for the difference lies in the observation that the stretch immediately prior to the predicted register error differs in the structure, forming either an uninterrupted β-strand (4x5p) or a β-bulge (Richardson *et al*., 1978) (4xo9): indeed this difference triggers the downstream register difference. Crucially, however, this surface-exposed stretch is at the crystal lattice interface in each of the structures, and the lattice differs since the crystals are in different space groups (Supp Fig 4b). In particular, hydrogen bonds made by the side chain of Arg60 in each form may help drive the local structural difference (Supp Fig 4c). Fold-switching proteins with more dramatic differences between alternative biologically relevant structures are considered below.

The second example involves structures of tick anticoagulant peptide at both high (1.62 Å; PDB 1dod; chain A) and low resolution (3.0 Å, PDB 1kig; chain I). They share a 1-residue register shift with respect to the AF2 model over residues 19-28. Although, unfortunately, no diffraction data are available, the crystal structures also share the same register with two NMR structures of the same protein (PDB 1tcp and 1tap) suggesting that here the AF2 model, and hence the error prediction, is either straightforwardly wrong or captures an alternative legitimate conformation. Notably, this protein contains three disulphide bonds and some literature suggests these may lead to difficulties for AF2 (Thornton *et al*., 2021; Wehrspan *et al*., 2022). It is also worth noting that the AF2 model can be fit to the crystal structures only rather poorly with a 2.89-3.03 rmsd on 59 Cα atoms.

The third example features structures of *Methylophilus methylotrophus* flavoprotein at high (1.60 Å; PDB 1o97; chain A) and low resolution (3.1 Å, PDB 1o96; chain Z). They share a 1-residue register shift with respect to the AF2 model over residues 196-201. Immediately prior to this region they each have a number of residues, three or six, that are not traced in the final structure: this relates to the lower local quality of the electron density in the domain linker region. Although not allowing for immediate and unambiguous interpretation, the electron density calculated using the higher resolution structure seemed to be more consistent with the AF2-preferred register (not shown). Indeed, the register matching the AF2 model is seen in structures deposited six years later by the same group e.g. PDB 3cls at 1.65 Å.

### 3.4 Local correlation values improve when predicted errors are corrected

Deposited models where a sequence register error was predicted were corrected as described in Section 2.5X.Y. Then, either the Density-fitness real-space correlation coefficient (RSCC) (MX cases) or the PHENIX map-model correlation coefficient (CC, cryo-EM cases) was calculated for the stretch of residues that were modified, and compared with the coefficient achieved with the set of residues present in the original deposition. Comparison of these local correlation values revealed that the CC improved in 4522 of 5599 cases (80%) of the cryo-EM cases and the RSCC improved in 3266 of 4059 cases (80%) of the MX cases (Figure 5).

**Figure 5.**
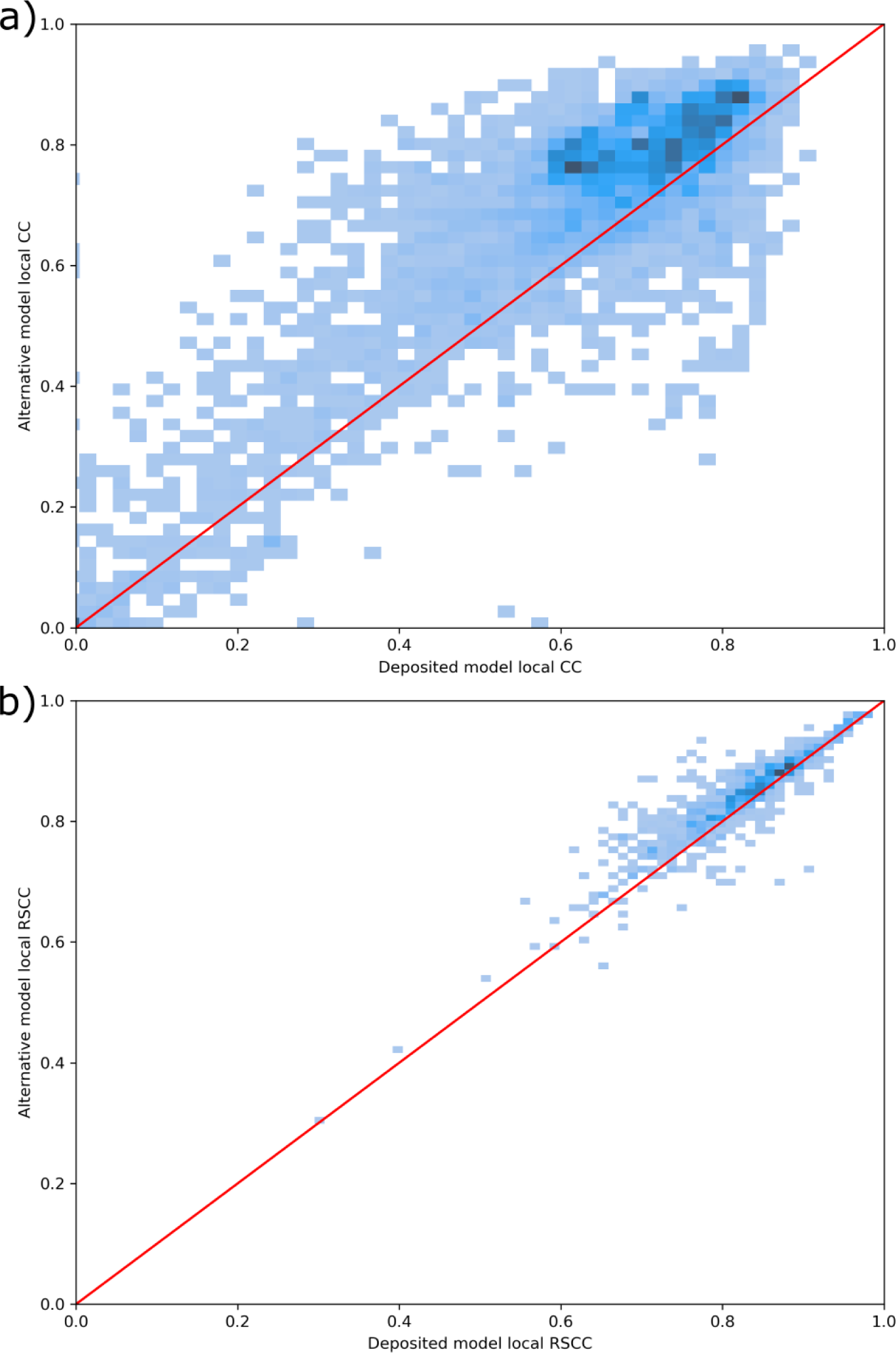
Density plots illustrate that local CC values improve for most cryoEM (A; PHENIX map-model correlation coefficient) and MX (B; Density-fitness real-space correlation coefficient) cases after correction of predicted errors.

### 3.5 Comparison with other validation tools

#### 3.5.1 New generation map-based validation with checkMySequence

In order to cross-validate the predicted errors that we found in PDB deposits, we used cMS to look for possible register errors within the structures in our dataset. cMS is based on map-model compatibility and hence is a completely independent approach. Since cMS is based on comparison of the model with the experimental data, only those models that had experimental data deposited could be cross-validated. Out of the 16,662 PDB entries that we analysed, 12,879 met this requirement. Furthermore, due to the underlying methods used within cMS, only register errors spanning at least 10 residues will be flagged. That means, out of the 4606 “unique” errors that we found in the dataset, only 1556 could be potentially found by cMS (33%).

A total of 672 errors were predicted by cMS among the 12,879 entries in the dataset that contained experimental data. Of these, only 90 were part of the set of 1556 errors also predicted by conkit-validate. A comparison of the resolution distributions of the deposited structures where an error was or wasn’t found using cMS is shown in Figure 6. The distribution of predicted errors extending to lower resolution cases in the set found by conkit-validate alone illustrates the valuable resolution independence of the program.

**Figure 6.**
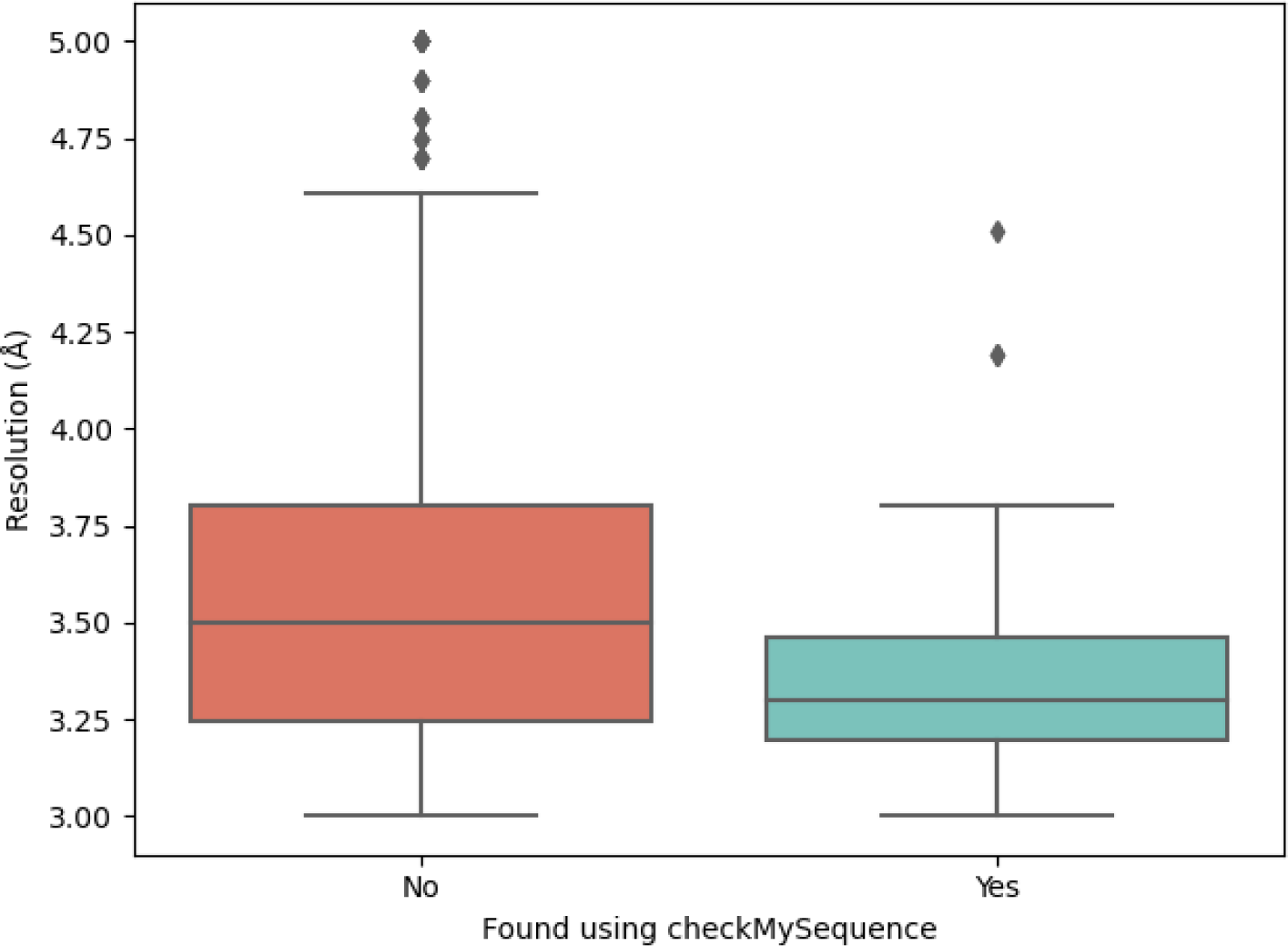
For those deposited structures where a possible sequence register was predicted by conkit-validate and had experimental data available, comparison of the distribution of the resolution at which they were solved and whether a possible register error was also predicted by checkMysequence.

Additionally, analysis of the structures where a sequence register error was flagged by cMS but not by conkit-validate revealed that 70% had been predicted to contain the same error by conkit-validate, but these errors were discarded by the filters described in section 2.3 and established in a previous study (Sánchez Rodríguez et al., 2022). This suggests that there is room for improvement in the filtering of results that we previously found necessary to reduce the incidence of false positive results (Sánchez Rodríguez et al., 2022) i.e. it has removed some true positives that were also discovered by cMS.

#### 3.5.2 Model geometry

It has been observed that register shifts may result in local concentrations of model geometry violations (Chojnowski, 2022, 2023) we evaluated fractions of various geometry errors identified using MolProbity (Ramachandran plot outliers, rotamer outliers, and Cβ deviations) in model-fragments with identified register shifts and random fragments of comparable lengths selected from the same models. We observed that the fraction of errors is indeed larger in regions with putative errors, but the difference is relatively small for all the validation criteria tested (Figure 7). This clearly supports our previous observation that register shifts may result in a locally increased number of validation outliers. However, it is clear that many regions with confident putative register errors identifiable with our new method lack geometric issues that alone would enable their identification.

**Figure 7.**
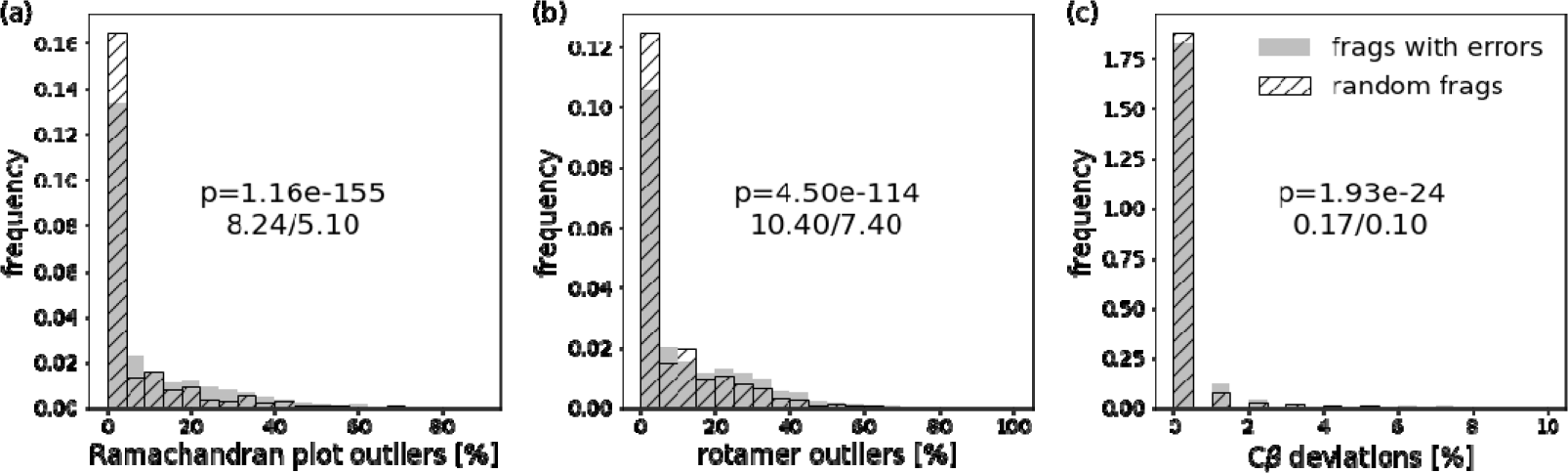
Comparison of distributions of (a) Ramachandran plot outliers (b) rotamer outliers and (c) Cβ deviations detected using Molprobity in model-fragments with plausible register shifts (grey bars) and random fragments (hatched bars). The p-values shown on the plots correspond to a two-sample t-test with an hypothesis that the expectation value for random fragments is smaller. The numbers below the p-values are mean fractions of outliers for fragments with errors and random fragments respectively.

#### 3.5.3 The composite cryo-EM validation measure DAQ-score

We further assessed the predicted register errors for the cryo-EM targets by comparing the average ΔCC score between the original and the alternative models against the average DAQ score over the predicted register error (Figure 8). Where DAQ score indicated a potential error (average DAQ AA < 0) we observed that ΔCC improved in 1699 of the 2136 cases (79.5%). Where no error was indicated by DAQ score (average DAQ AA > 0) we observed that ΔCC improved in 2525 of the 3758 cases (67.2%). It should be noted, in a large number of these cases, the average DAQ score was very close to zero, indicating that the map lacked distinctive map features that would allow DAQ’s neural network to make a decisive call. This helps to highlight the importance of our map-independent method when looking for potential errors.

**Figure 8.**
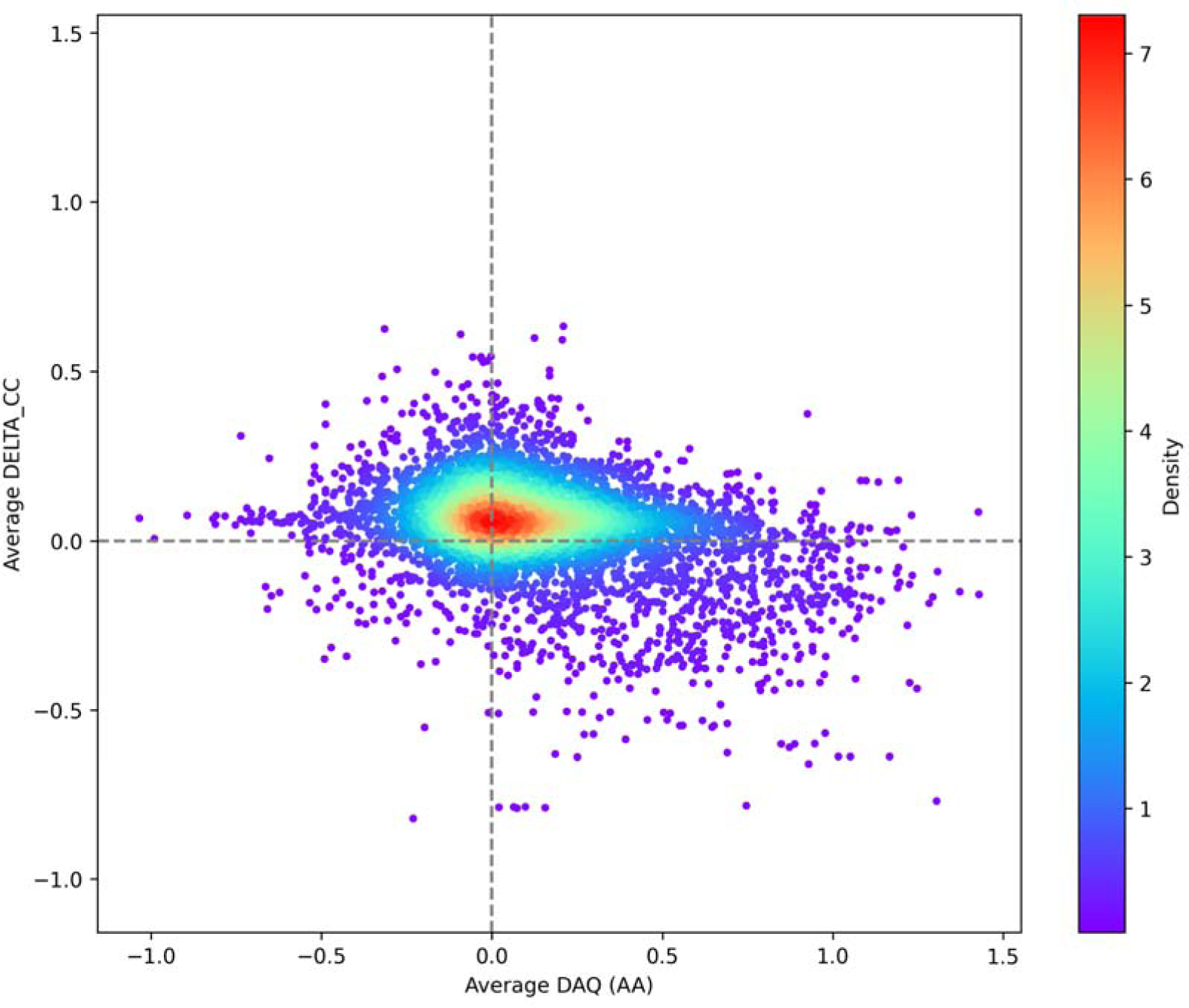
A scatter diagram showing average delta in PHENIX map-model CC before and after correction, versus the average DAQ score over the predicted error. The points in the plot are coloured by density where blue represents a low density of points and red represents a high density of points.

### 3.6 Selected examples

#### 3.6.1 cryo-EM examples

Fig 9 illustrates two representative predicted errors discovered by conkit-validate. The first, also flagged by cMS (A), indicates a predicted 18-residues error in chain SB (note that we use author chain IDs throughout) of the Escherichia coli 30S ribosomal protein S2 of a ribosome structure, PDB 3j9z determined to a reported resolution of 3.6 Å. The second, found solely by conkit-validate (B) illustrates a predicted 28-residue error in chain BJ of the rabbit ribosome structure, PDB 6gz3, also at 3.6 Å. Notably, the regions in question, when analysed by CryoNet (Dai et al., 2023) appear to have local map resolution significantly worse than the overall quoted values and certainly worse than 4 Å (Supp Fig 5). In each case, bulky, aromatic side chains provide visual confirmation that the new register is correct. Indeed, the local map-model CC for the region in question improves dramatically after correction of the error as suggested by conkit-validate: for 3j9z it improves from 0.29 to 0.71, while the values before and after correction for 6gz3 are 0.51 and 0.82.

**Figure 9.**
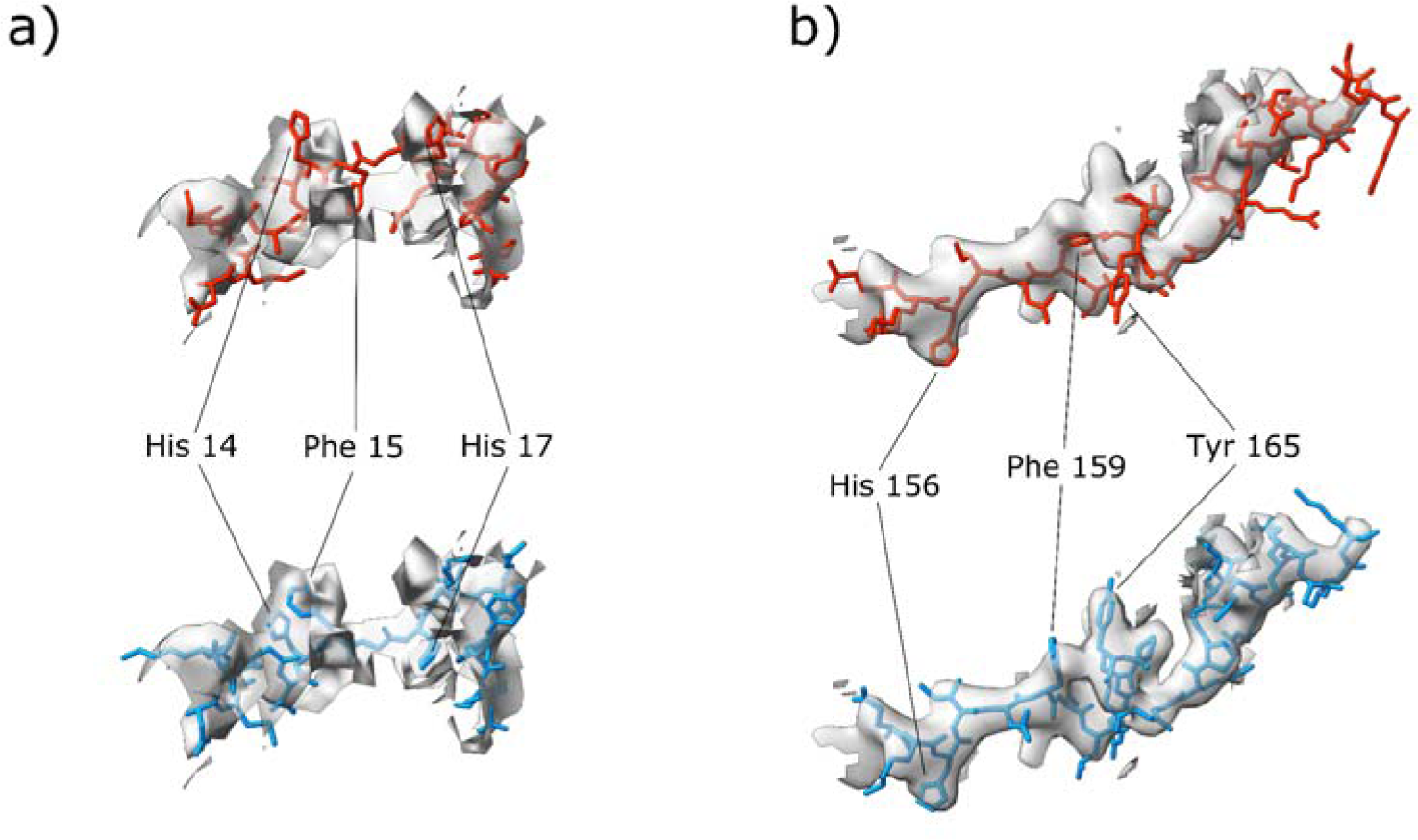
Detailed views of portions of deposited cryo-EM structures in which a possible sequence-register error was detected using conkit-validate. A) The error corresponds to PDB entry 3j9z chain SB residues 7–24 and residues 14, 15 and 17 have been highlighted. B) The error corresponds to PDB entry 6gz3 chain BJ residues 153–180 and residues 156, 159 and 165 have been highlighted. In each case, mask of 2.5 Å around the model was applied; and the original deposition is coloured red with the structure corrected with the sequence register suggested by conkit-validate in blue. The density map for the deposited structure is represented as a transparent grey surface with the contour level set to 0.035 in (A) and 2.2 in (B).

#### 3.6.2 MX examples

Fig 10 shows representative examples of predicted register errors in crystal structures, before and after correction. The predicted errors are found in a bacterial ion channel determined to 3.3 Å resolution (A), a human coagulation factor complex at 4.19 Å (B) and a bacterial carbonic anhydrase at 3.17 Å (C). The first two were also detected by cMS, the last not. The case in (A) is the same error highlighted in a cryo-EM structure in our previous paper (Sánchez Rodríguez et al., 2022), illustrating the map-agnostic nature of our method.

**Figure 10.**
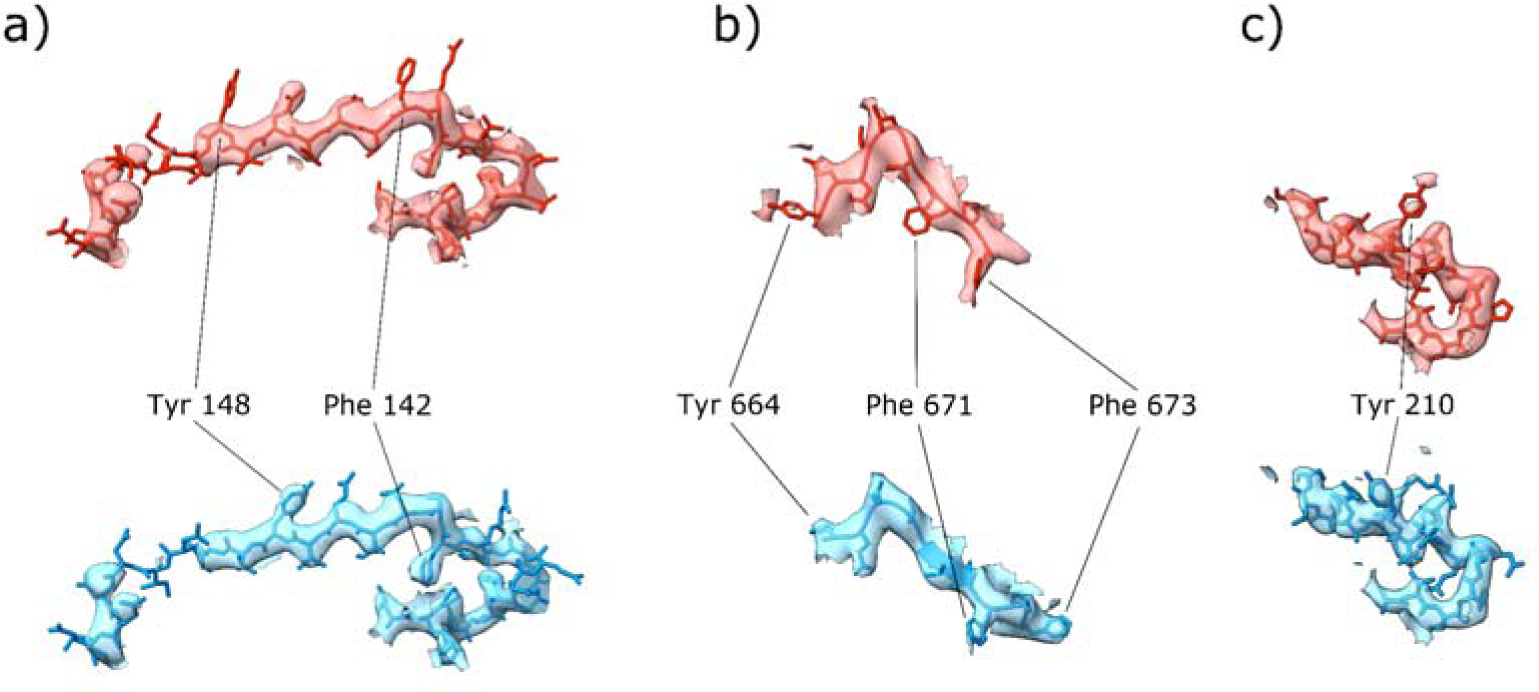
Detailed views of portions of deposited MX structures in which a possible sequence-register error was detected using conkit-validate. A) The error corresponds to PDB entry 2yks chain A residues 130–157 and residues 142 and 148 have been highlighted. B) The error corresponds to PDB entry 5k8d chain A residues 664–673 and residues 664, 671 and 673 have been highlighted. C) The error corresponds to PDB entry 6yl7 chain A residues 199–219 and residue 210 have been highlighted. In each case, mask of 2.4 Å around the model was applied. The maps are 2*F* _o_ − *F* _c_ maps coloured at the same contour level with the original deposition in red and the structure corrected with the sequence register suggested by conkit-validate in blue. The contour levels are 0.207, 0.154 and 0.24 in A) to C).

#### 3.6.3 Fold-switching

Our method depends on spotting mismatches between a deposited structure and the structure implied by deep learning-based prediction of inter-residue distances and contacts. It will therefore potentially struggle with the small proportion of proteins, known as fold-switchers, which have alternative, distinctly different, but equally biologically valid folds. We find one such example in our set, human calcineurin (PDB 5c1v, determined by X-ray crystallography to 3.35 Å) which undergoes a significant rearrangement of C-terminal β-strands as a result of reversible cis-trans isomerisation of a proline residue (Guasch *et al*., 2015). In the structure chain A, with trans-Pro 309 adopted the previously universal structure, whereas cis-Pro 309 in chain B led to a novel conformation in the C-terminal region and, in particular, a different sequence register of two β-strands compared to chain A (Fig 11). Electron density provides good support for the novel conformation (Fig 11b) yet residues 288-340 are flagged as a potential error by conkit-validate. The attempted ‘correction’ of register leads to a confirmation resembling that of chain A which has a poorer fit to the density (Fig 11d) demonstrating that this case is a false positive error prediction.

**Figure 11.**
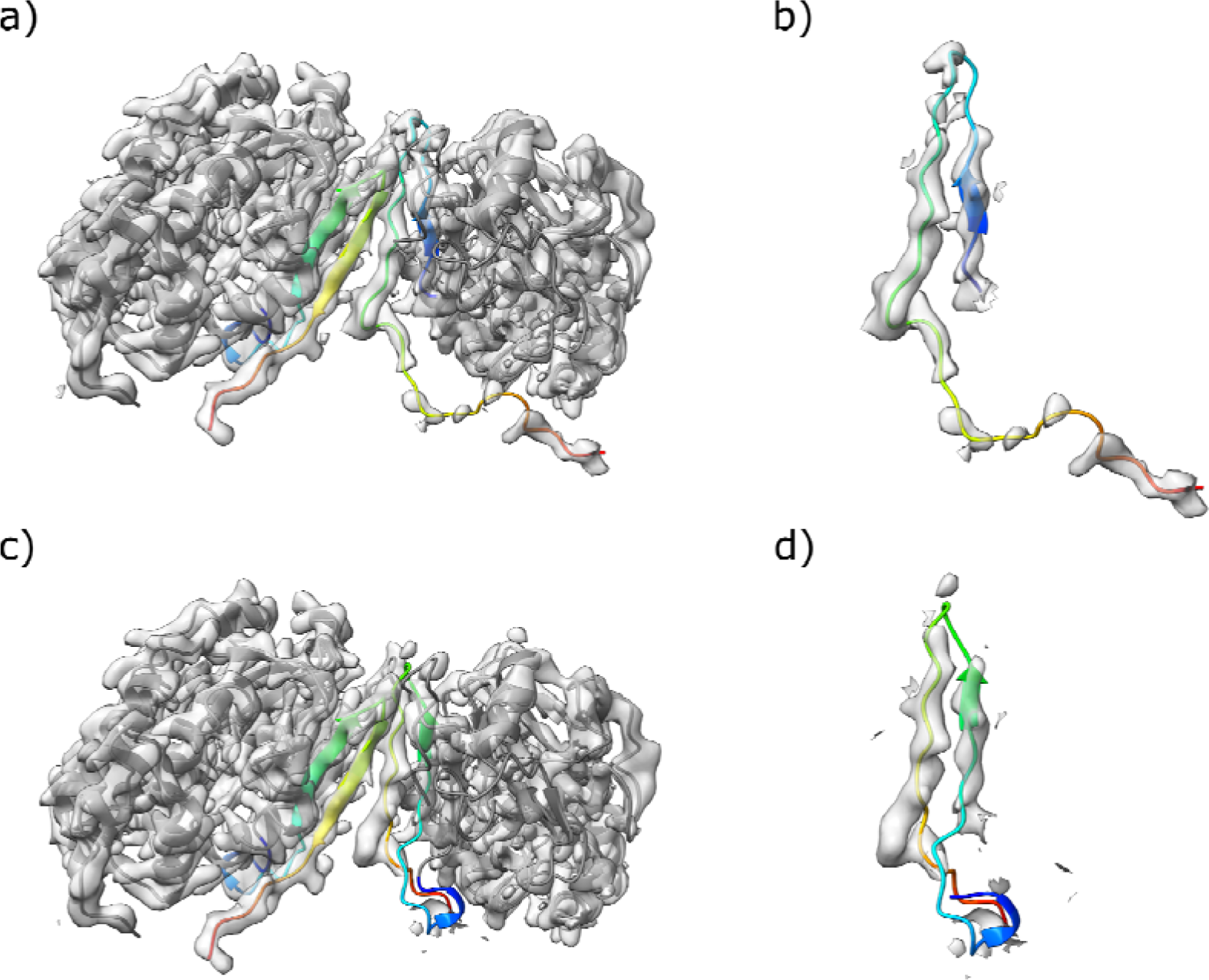
Detailed view of the section of the deposited model where conkit-validate erroneously identified a potential sequence-register error in human calcineurin (PDB 5c1v). a) the deposited structure is shown in grey with the fold-switching region (residues 309-340) shown in a rainbow spectrum where blue indicates its N-terminal part and red indicates the C-terminus. The density map for the deposited structure is represented as a transparent grey surface and the contour level was set to 0.25. b) a close up of the fold-switching region in chain B of the deposited structure and the associated density. c) the automated attempt at fixing the potential sequence-register error identified by conkit-validate. d) a close up of the fold-switching region in chain B of the attempted fix.

## 4. Conclusions

We have presented a PDB-wide analysis of deposited structures determined to between 3 and 5Å resolution using our new validation protocol. It is based on deep-learning derived predictions of inter-residue distances and contacts, but the availability of an associated 3D-structure prediction provides extra confidence by enabling filtering out of some false positives (Sánchez Rodríguez *et al*., 2022). Potential errors are flagged as disparities between the observed residue distances and contacts and those predicted by the latest generation of deep learning-based tools, most notably AlphaFold 2 (Jumper *et al*., 2021). Although general in its ability to detect errors, our method is particularly effective when applied, as here, to detection of register errors (Sánchez Rodríguez *et al*., 2022). Importantly, our method is orthogonal to current validation metrics, providing a valuable further tool for detection of errors in protein structure. Also notably, it is independent of the experimental map, be it cryo-EM or crystallographic, and hence its performance is unaffected by (local) resolution.

We found that around one in six PDB deposits determined to 3-5 Å resolution contained a predicted register error, although this amounts to only 2.3% of residues. Given the benchmarking previously carried out, the implementation of filters to catch false positives, and the observable improvements in map-model compatibility after correction (Fig xx), we are confident that these represent high-confidence predictions. The large numbers involved allow trends to be analysed: unsurprisingly, entries in the higher-resolution part of the studied range are less likely to contain a predicted error than lower-resolution deposits. We find that cryo-EM structures are more likely to contain a predicted error than crystallographic structures at the same quoted resolution, but it must be borne in mind that cryo-EM maps, especially, often contain regions where resolution is much poorer than the headline value. In addition, the gap of error frequency between the two methods seems to be narrowing recently.

While powerful, our method has limitations. Most obviously, it depends on the quality of the distance and contact predictions available. In most cases, the quality is very high, as illustrated by the superb accuracy achieved by methods such as AlphaFold 2 (Jumper *et al*., 2021; Pereira *et al*., 2021; Simpkin *et al*., 2023). However, recent analysis demonstrates that contemporary modelling methods of different kinds still consistently struggle with targets that are singletons or for which very few homologues can be found in databases. In addition, small size and high a-helical content may be additional aggravating factors (Simpkin *et al*., 2023). In such cases, where accurate modelling is not possible then the corresponding distance and contact predictions will also be lower quality and hence, potentially, not suitable for use in validation. In this regard, we have also presented an example where the presence of disulphide bonds in the target may have resulted in an unusually low quality AF2 prediction. As we have also shown, our method can also misbehave in rare cases of fold-switching proteins, flagging a potential error in the part of the structure that has two biologically valid structures. A similar situation will arise in very unusual cases of homologous proteins adopting entirely different structures (Schierholz *et al*., 2024). More positively, such a misprediction in regions of confident structure building could potentially be interpreted as a sign of a potentially interesting fold-switching region, or a different kind of biologically relevant structural ambivalence. It may also be that future work looking at AF2 predicted distance probabilities in more detail e.g. (Brown *et al*., 2024) can flag such regions directly, thereby avoiding these rare mispredictions.

In summary, our method, which is map-independent and orthogonal to prevalent validation software, effectively pinpoints register errors in large numbers of mid-resolution PDB entries, illustrating the challenges facing even diligent and expert structural biologists when working on this kind of target. Notably, and unlike other validation methods, ours provides a suggested correction to the register for putative errors. Finally, with simple adaptation to use Cα rather than Cβ atoms for definition of contacts, we expect our method could answer the call from the PDB cryo-EM Data Archiving and Validation Group recently (Kleywegt *et al*., 2024) for methods to validate Cα-only structures. We hope our method’s availability through CCP4 (Agirre *et al*., 2023) will contribute to detecting errors before deposition in the PDB.

## Acknowledgements

The authors thank Robbie Joosten for useful discussions.

## Data availability

The set of predicted register errors are available in Supp. Table 1.

conkit-validate is distributed with the CCP4 software suite (Agirre *et al*., 2023).

## References

Afonine, P. V., Sobolev, O. V., Moriarty, N. W., Terwilliger, T. C. & Adams, P. D. (2023). Acta Crystallogr D Struct Biol 10.1107/S2059798323005077.

Agirre, J., Atanasova, M., Bagdonas, H., Ballard, C. B., Baslé, A., Beilsten-Edmands, J., Borges, R. J., Brown, D. G., Burgos-Mármol, J. J., Berrisford, J. M., Bond, P. S., Caballero, I., Catapano, L., Chojnowski, G., Cook, A. G., Cowtan, K. D., Croll, T. I., Debreczeni, J. É., Devenish, N. E., Dodson, E. J., Drevon, T. R., Emsley, P., Evans, G., Evans, P. R., Fando, M., Foadi, J., Fuentes-Montero, L., Garman, E. F., Gerstel, M., Gildea, R. J., Hatti, K., Hekkelman, M. L., Heuser, P., Hoh, S. W., Hough, M. A., Jenkins, H. T., Jiménez, E., Joosten, R. P., Keegan, R. M., Keep, N., Krissinel, E. B., Kolenko, P., Kovalevskiy, O., Lamzin, V. S., Lawson, D. M., Lebedev, A. A., Leslie, A. G. W., Lohkamp, B., Long, F., Malý, M., McCoy, A. J., McNicholas, S. J., Medina, A., Millán, C., Murray, J. W., Murshudov, G. N., Nicholls, R. A., Noble, M. E. M., Oeffner, R., Pannu, N. S., Parkhurst, J. M., Pearce, N., Pereira, J., Perrakis, A., Powell, H. R., Read, R. J., Rigden, D. J., Rochira, W., Sammito, M., Sánchez Rodríguez, F., Sheldrick, G. M., Shelley, K. L., Simkovic, F., Simpkin, A. J., Skubak, P., Sobolev, E., Steiner, R. A., Stevenson, K., Tews, I., Thomas, J. M. H., Thorn, A., Valls, J. T., Uski, V., Usón, I., Vagin, A., Velankar, S., Vollmar, M., Walden, H., Waterman, D., Wilson, K. S., Winn, M. D., Winter, G., Wojdyr, M. & Yamashita, K. (2023). Acta Crystallogr D Struct Biol 79, 449–461.

Barad, B. A., Echols, N., Wang, R. Y.-R., Cheng, Y., DiMaio, F., Adams, P. D. & Fraser, J. S. (2015). Nat. Methods 12, 943–946.

Brown, B. P., Stein, R. A., Meiler, J. & Mchaourab, H. S. (2024). J. Chem. Theory Comput. 10.1021/acs.jctc.3c01081.

Callaway, E. (2020). Nature 578, 201.

Casañal, A., Lohkamp, B. & Emsley, P. (2020). Protein Sci. 29, 1069–1078.

Chojnowski, G. (2022). Acta Crystallogr D Struct Biol 78, 806–816.

Chojnowski, G. (2023). Acta Crystallogr D Struct Biol 79, 559–568.

Colovos, C. & Yeates, T. O. (1993). Protein Sci. 2, 1511–1519.

Croll, T. I. (2018). Acta Crystallographica Section D: Structural Biology 74, 519–530.

Dai, M., Dong, Z., Xu, K. & Zhang, Q. C. (2023). J. Mol. Biol. 435, 168059.

Davis, I. W., Leaver-Fay, A., Chen, V. B., Block, J. N., Kapral, G. J., Wang, X., Murray, L. W., Arendall, W. B., 3rd, Snoeyink, J., Richardson, J. S. & Richardson, D. C. (2007). Nucleic Acids Res. 35, W375–W383.

Eddy, S. R. (2011). PLoS Comput. Biol. 7, e1002195.

Emsley, P. & Cowtan, K. (2004). Acta Crystallogr. D Biol. Crystallogr. 60, 2126–2132.

Gao, Y., Thorn, V. & Thorn, A. (2023). Acta Crystallogr D Struct Biol 79, 206–211.

GitHub - PDB-REDO/density-fitness: Application to calculate the density statistics (RSR, SRSR, RSCCS, EDIAm and OPIA) for x-ray structures GitHub, https://github.com/PDB-REDO/density-fitness.

Guasch, A., Aranguren-Ibáñez, Á., Pérez-Luque, R., Aparicio, D., Martínez-Høyer, S., Mulero, M. C., Serrano-Candelas, E., Pérez-Riba, M. & Fita, I. (2015). PLoS One 10, e0134569.

Joosten, R. P., Joosten, K., Murshudov, G. N. & Perrakis, A. (2012). Acta Crystallogr. D Biol. Crystallogr. 68, 484–496.

Joseph, A. P., Malhotra, S., Burnley, T., Wood, C., Clare, D. K., Winn, M. & Topf, M. (2016). Methods 100, 42–49.

Jumper, J., Evans, R., Pritzel, A., Green, T., Figurnov, M., Ronneberger, O., Tunyasuvunakool, K., Bates, R., Žídek, A., Potapenko, A., Bridgland, A., Meyer, C., Kohl, S. A. A., Ballard, A. J., Cowie, A., Romera-Paredes, B., Nikolov, S., Jain, R., Adler, J., Back, T., Petersen, S., Reiman, D., Clancy, E., Zielinski, M., Steinegger, M., Pacholska, M., Berghammer, T., Bodenstein, S., Silver, D., Vinyals, O., Senior, A. W., Kavukcuoglu, K., Kohli, P. & Hassabis, D. (2021). Nature 596, 583–589.

Kleywegt, G. J., Adams, P. D., Butcher, S. J., Lawson, C. L., Rohou, A., Rosenthal, P. B., Subramaniam, S., Topf, M., Abbott, S., Baldwin, P. R., Berrisford, J. M., Bricogne, G., Choudhary, P., Croll, T. I., Danev, R., Ganesan, S. J., Grant, T., Gutmanas, A., Henderson, R., Heymann, J. B., Huiskonen, J. T., Istrate, A., Kato, T., Lander, G. C., Lok, S. M., Ludtke, S. J., Murshudov, G. N., Pye, R., Pintilie, G. D., Richardson, J. S., Sachse, C., Salih, O., Scheres, S. H. W., Schroeder, G. F., Sorzano, C. O. S., Stagg, S. M., Wang, Z., Warshamanage, R., Westbrook, J. D., Winn, M. D., Young, J. Y., Burley, S. K., Hoch, J. C., Kurisu, G., Morris, K., Patwardhan, A. & Velankar, S. (2024). IUCrJ 11, 140–151.

Krissinel, E. (2012). J Mol Biochem 1, 76–85.

Laskowski, R. A., MacArthur, M. W., Moss, D. S. & Thornton, J. M. (1993). J. Appl. Crystallogr. 26, 283–291.

Liebschner, D., Afonine, P. V., Baker, M. L., Bunkóczi, G., Chen, V. B., Croll, T. I., Hintze, B., Hung, L.-W., Jain, S., McCoy, A. J., Moriarty, N. W., Oeffner, R. D., Poon, B. K., Prisant, M. G., Read, R. J., Richardson, J. S., Richardson, D. C., Sammito, Sobolev, O. V., Stockwell, D. H., Terwilliger, T. C., Urzhumtsev, A. G., Videau, L. L., Williams, C. J. & Adams, P. D. (2019). Acta Crystallographica Section D: Structural Biology 75, 861–877.

Lüthy, R., Bowie, J. U. & Eisenberg, D. (1992). Nature 356, 83–85.

Mirdita, M., Steinegger, M. & Söding, J. (2019). Bioinformatics 35, 2856–2858.

Murshudov, G. N., Skubák, P., Lebedev, A. A., Pannu, N. S., Steiner, R. A., Nicholls, R. A., Winn, Long, F. & Vagin, A. A. (2011). Acta Crystallogr. D Biol. Crystallogr. 67, 355–367.

Nakamura, T., Wang, X., Terashi, G. & Kihara, D. (2023). Nat. Methods 20, 775–776.

Ovchinnikov, S., Park, H., Varghese, N., Huang, P.-S., Pavlopoulos, G. A., Kim, D. E., Kamisetty, H., Kyrpides, N. C. & Baker, D. (2017). Science 355, 294–298.

Pereira, J., Simpkin, A. J., Hartmann, M. D., Rigden, D. J., Keegan, R. M. & Lupas, A. N. (2021). Proteins 89, 1687–1699.

Pintilie, G., Zhang, K., Su, Z., Li, S., Schmid, M. F. & Chiu, W. (2020). Nat. Methods 17, 328–334.

Prisant, M. G., Williams, C. J., Chen, V. B., Richardson, J. S. & Richardson, D. C. (2020). Protein Sci. 29, 315–329.

Reggiano, G., Lugmayr, W., Farrell, D., Marlovits, T. C. & DiMaio, F. (2023). Structure 31, 860–869.e4.

Richardson, J. S., Getzoff, E. D. & Richardson, D. C. (1978). Proc. Natl. Acad. Sci. U. S. A. 75, 2574–2578.

Sali, A., Berman, H. M., Schwede, T., Trewhella, J., Kleywegt, G., Burley, S. K., Markley, J., Nakamura, H., Adams, P., Bonvin, A. M. J. J., Chiu, W., Peraro, M. D., Di Maio, F., Ferrin, T. E., Grünewald, K., Gutmanas, A., Henderson, R., Hummer, G., Iwasaki, K., Johnson, G., Lawson, C. L., Meiler, J., Marti-Renom, M. A., Montelione, G. T., Nilges, M., Nussinov, R., Patwardhan, A., Rappsilber, J., Read, R. J., Saibil, H., Schröder, G. F., Schwieters, C. D., Seidel, C. A. M., Svergun, D., Topf, M., Ulrich, E. L., Velankar, S. & Westbrook, J. D. (2015). Structure 23, 1156–1167.

Sánchez Rodríguez, F., Chojnowski, G., Keegan, R. M. & Rigden, D. J. (2022). Acta Crystallographica Section D: Structural Biology 78, 1412–1427.

Schierholz, L., Brown, C. R., Helena-Bueno, K., Uversky, V. N., Hirt, R. P., Barandun, J. & Melnikov, S. V. (2024). Mol. Biol. Evol. 41, 10.1093/molbev/msad254.

Simpkin, A. J., Mesdaghi, S., Sánchez Rodríguez, F., Elliott, L., Murphy, D. L., Kryshtafovych, A., Keegan, R. M. & Rigden, D. J. (2023). Proteins 10.1002/prot.26593.

Sippl, M. J. (1993). Proteins 17, 355–362.

Terashi, G., Wang, X., Maddhuri Venkata Subramaniya, S. R., Tesmer, J. J. G. & Kihara, D. (2022). Nat. Methods 19, 1116–1125.

Thornton, J. M., Laskowski, R. A. & Borkakoti, N. (2021). Nat. Med. 27, 1666–1669.

Vriend, G (1990). J. Mol. Graph. 8, 52–56.

Vriend, G. & Sander, C. (1993). J. Appl. Crystallogr. 26, 47–60.

Wehrspan, Z. J., McDonnell, R. T. & Elcock, A. H. (2022). J. Mol. Biol. 434, 167377.

wwPDB consortium (2019). Nucleic Acids Res. 47, D520–D528.

