## Supplementary Material for "Using deep learning predictions reveals a large number of register errors in PDB deposits"

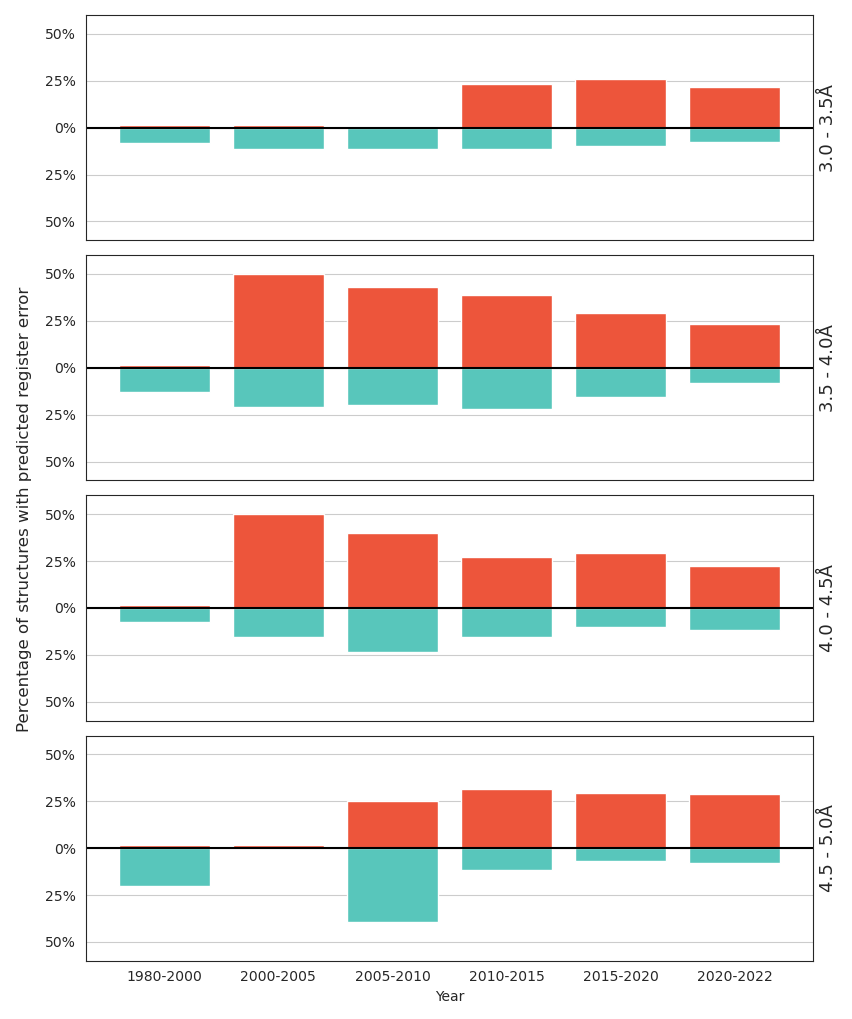

Supplementary Figure 1. Ratio of structures containing a predicted register error across different resolutions and years. Ratios for structures determined using MX have been coloured in blue while those determined with cryo-EM in red. Each row contains data for a given resolution bin. Horizontal axis indicates the binned year of deposition and the vertical axis the percentage of structures with a predicted error for that particular year and resolution bin.

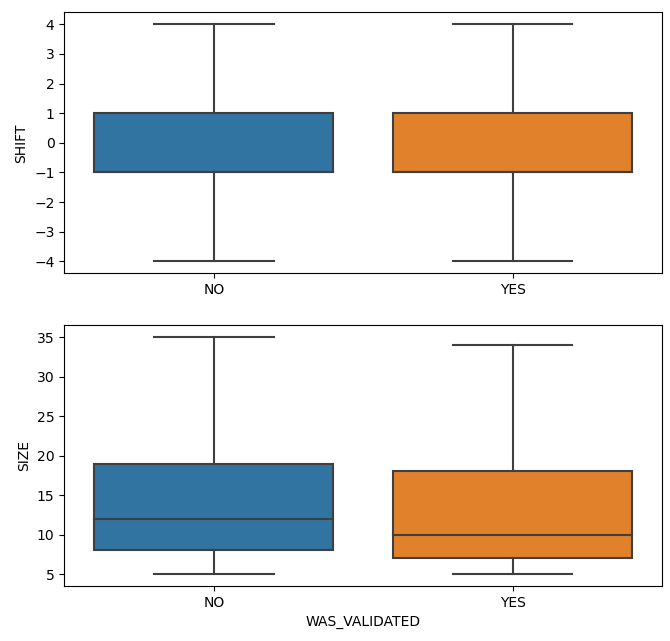

Supplementary Figure 2 Distribution of the shift required for correction of the predicted error (above) and number of residues affected by the predicted error (below) for the full set of 4606 errors (left) and the subset of 147 structure (403 errors) that could be validated by comparison to high-resolution MX structures.

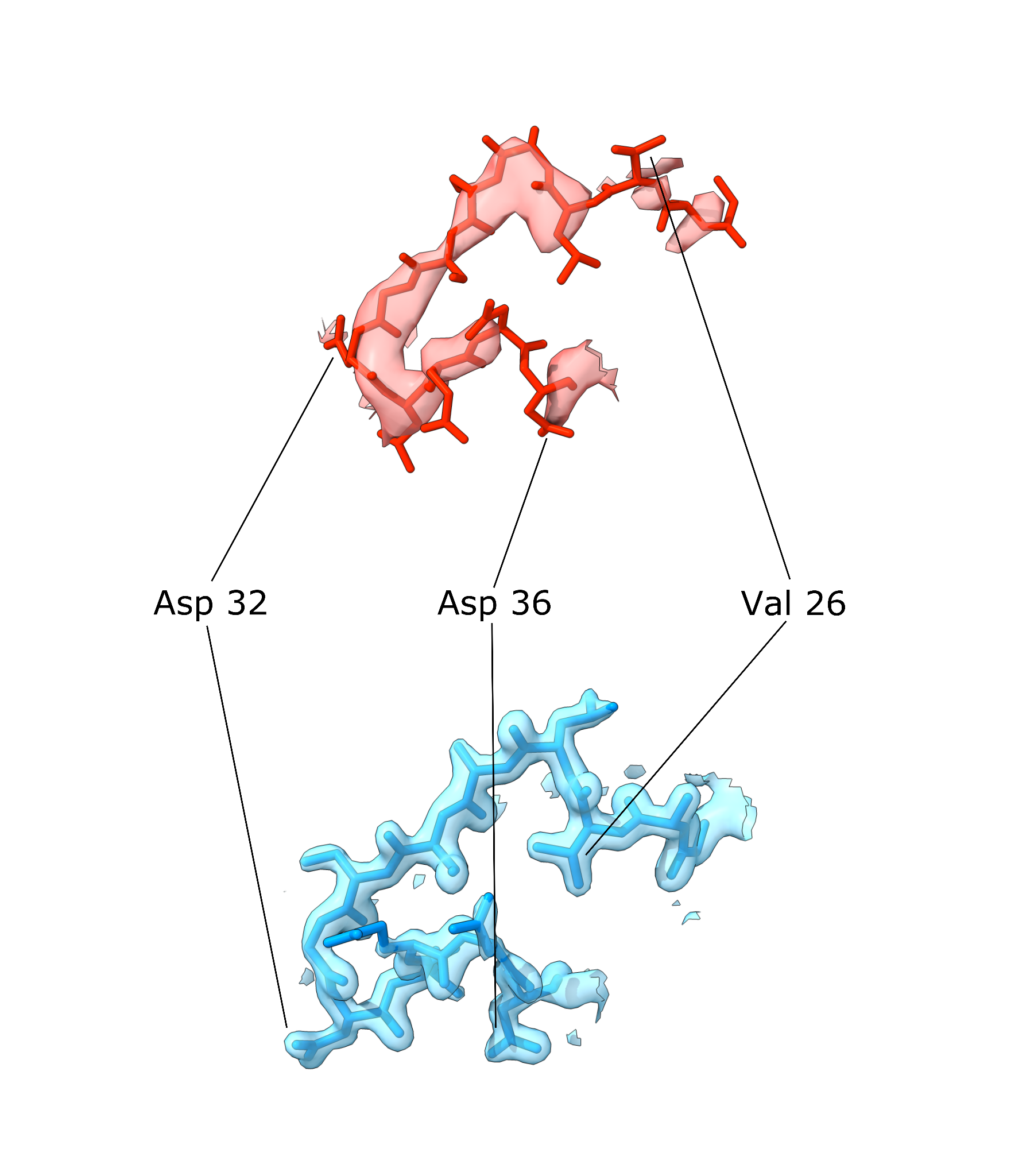

Supplementary Figure 3. Detailed view of a possible sequence-register error detected using conkit-validate in a structure solved at 3.01 Å using MX (red) (Ragulator complex protein LAMTOR4, PDB entry 5YK3, chain I, residues 24-36). Another structure for the same protein was solved using MX at 1.42Å (blue) which supports the alternative sequence register proposed by conkit-validate (PDB entry 6B9X, chain D, residues 24-36). In each case, mask of 2.5 Å around the model was applied with the contour level set to 0.27 for 5YK3 and 0.66 for 6B9X.

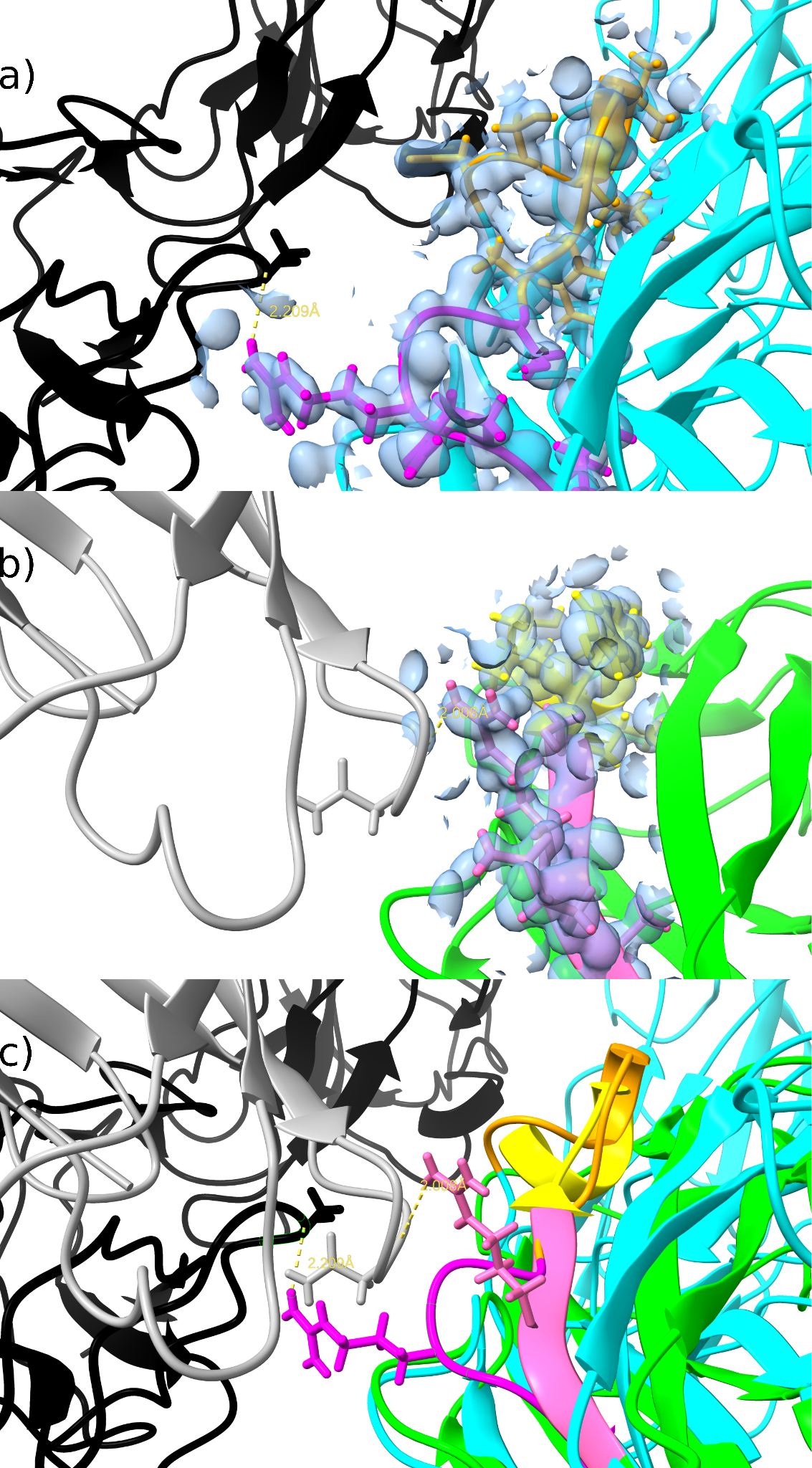

Supplementary Figure 4. Analysis and comparison of FimH structures. Electron density at contour levels 0.7/0.5 is shown in blue for residues 58-70 of structures with PDB codes 4xo9 (a) and 4x5p (b). c) A superposition of the two - 4xo9 and 4x5p as cyan/green with the predicted error region is shown in orange/yellow, the preceding region discussed in the text in magenta/pink - including Arg60 in sticks (with alternate conformations in one structure), crystal symmetry mates in black/grey, and potential hydrogen bonds as dotted yellow lines labelled with their length in A.

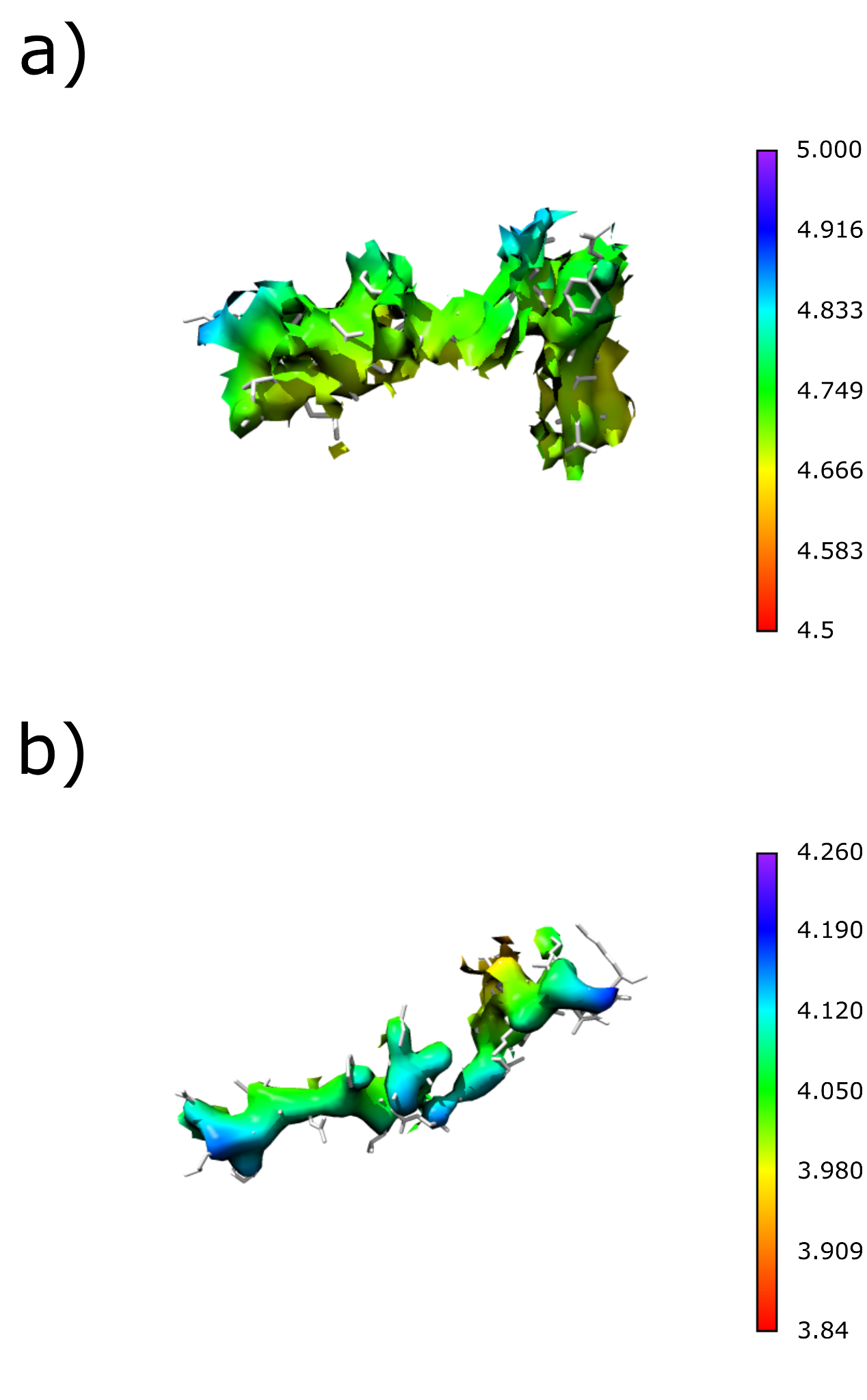

Supplementary Figure 5. The local resolutions, as reported by CryoNet, in the regions relating to conkit-validate predicted errors are seen to be significantly lower than the overall quoted resolution for a) 3j9z (3.6 Å) and b) 6gz3 (3.6 Å).

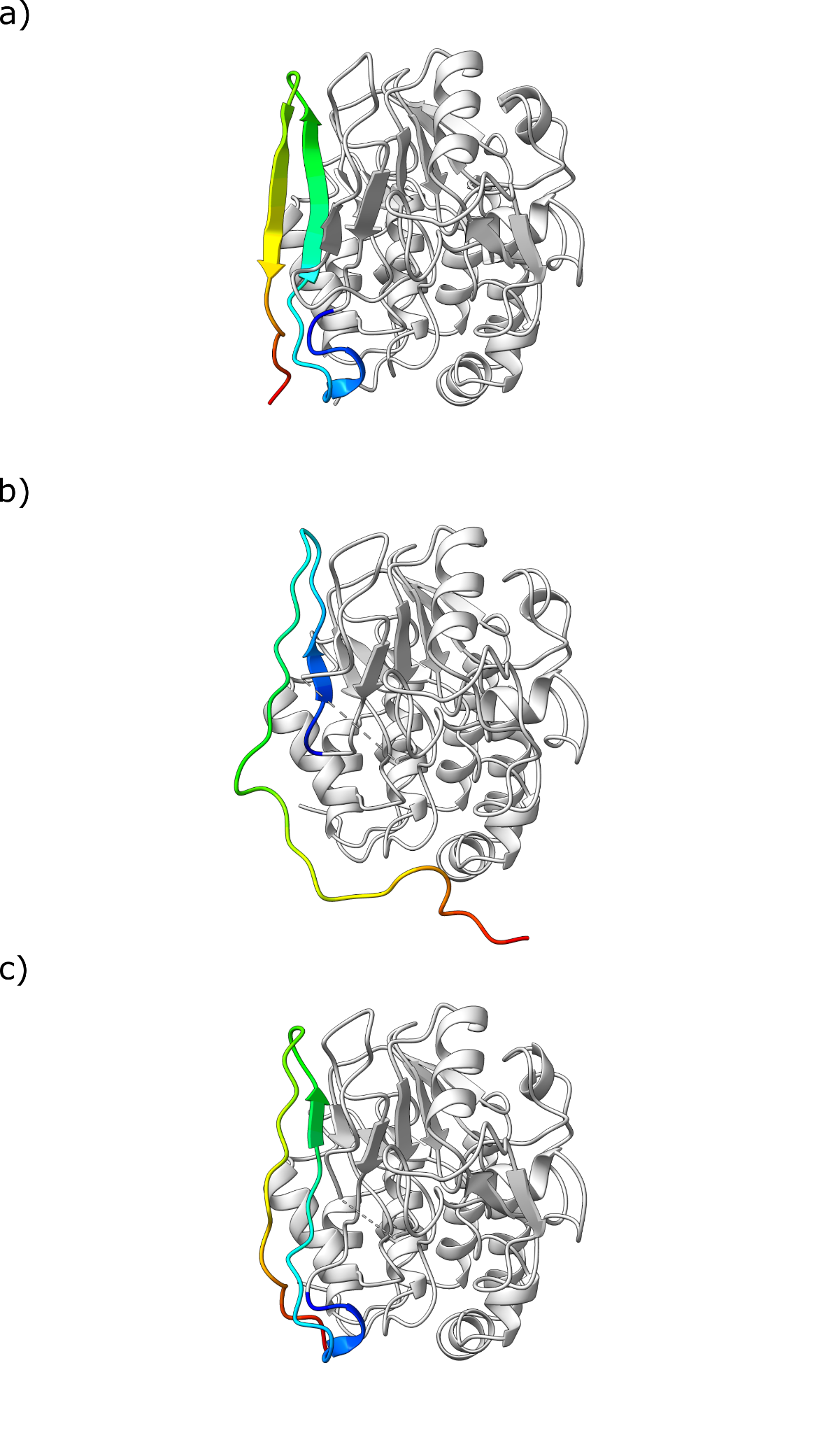

Supplementary Figure 6. Detailed view of the section of the deposited model where conkit-validate misidentified a fold-switching region as a potential sequence-register error in human calcineurin (PDB 5c1v where a) shows chain A of the deposited structure, b) shows chain B of the deposited structure and c) shows chain B with the attempted fix. In all cases, the majority of the model is shown in grey with the fold-switching region (residues 309-340) shown in a rainbow spectrum where blue indicates its N-terminal part and red indicates the C-terminus.

Supplementary Table 1. A csv file containing all the predicted errors found.

| Key | Description |
| --- | --- |
| PDB_CHAIN_ID | PDB code and the chain name where the error was predicted. |
| METHOD | Experimental method used to determine the structure. |
| RESOLUTION | Resolution of the deposited data. |
| YEAR | Deposition year. |
| CLUSTER_ID | Cluster identification number. Clusters consist of structures sharing 100% sequence similarity. |
| PLDDT | The average PLDDT for the region with the predicted error. |
| RESIDUE_RANGE | The residue range with the predicted error. |
| SIZE | The number of residues affected by the predicted error. |
| SHIFT | Average predicted shift for the predicted error. |
| GESAMT_QSCORE | The Q-Score reported by GESAMT for the alignment between AF2 predicted model and the deposited structure. |
| GESAMT_RMSD | The RMSD reported by GESAMT for the alignment between AF2 predicted model and deposited structure. |
| GESAMT_N_ALLIGNED | The number of residues aligned using GESAMT between AF2 predicted model and deposited structure (as a fraction of 1.0). |
| N_CONTACTS_MEAN | The mean number of contacts available for the contact map alignment in the region with the predicted error. |
| N_CONTACTS_MEDIAN | The median number of contacts available for the contact map alignment in the region with the predicted error. |
| N_CONTACTS_SUM | The total sum of the number of contacts available for the contact map alignment in the region with the predicted error. |
